# The *Vitis vinifera* receptor VvLYK6 negatively regulates chitin-triggered immune responses and promotes fungal infections

**DOI:** 10.1101/2025.06.06.658283

**Authors:** Jérémy Villette, Tania Marzari, David Landry, Thibault Roudaire, Agnès Klinguer, Nathalie Leborgne-Castel, Céline Vicedo, Virginie Gasciolli, Cécile Pouzet, Benoît Lefebvre, Marie-Claire Héloir, Benoit Poinssot

## Abstract

*Botrytis cinerea* is recognized as one of the most damaging fungal pathogens affecting grapevine (*Vitis vinifera*), directly impacting both grape yield and wine quality. Identifying new genes involved in the interaction between *V. vinifera* and *B. cinerea* appears to be a promising strategy for enhancing grapevine resistance in future breeding programs.

During pathogen infection, plasma membrane-localized pattern recognition receptors (PRRs) are responsible for detecting conserved microbe-associated molecular patterns (MAMPs). Among PRRs, members of the LysM receptor-like kinase family are well known to mediate the recognition of fungal MAMPs and triggering plant immune signaling pathways. Interestingly, a novel member of this receptor family, named VvLYK6, was identified in grapevine as the most highly upregulated during *B. cinerea* infection.

To investigate the role of VvLYK6 in plant immunity, we conducted overexpression studies in *Arabidopsis thaliana* and grapevine cell suspensions. Overexpression of *VvLYK6* led to a reduction in chitin-induced MAPK activation, decreased expression of defense-related genes, reduced callose deposition, and increased plant susceptibility to fungal pathogens in *A. thaliana*.

Based on these findings, we conclude that VvLYK6 acts as a negatively regulator of chitin-triggered immune responses, suggesting its potential role as a susceptibility gene during fungal infections.

## Introduction

Grapevine (*Vitis vinifera*) is continuously exposed to intense pest pressure, including viruses, insects, oomycetes such as downy mildew (*Plasmopara viticola*) and fungi such as powdery mildew (*Erysiphe necator*) or grey mold (*Botrytis cinerea*). The current strategy to mitigate the impact of these diseases relies heavily on chemical treatments. However, these treatments pose risks to both the environment and human health, prompting the development of alternative approaches to disease management in vineyards (Pathak et al., 2022). One such approach involves the identification of resistance (*R*) and susceptibility (*S*) genes (Qiu et al., 2015; Zaidi et al., 2018). The introgression of *R* genes into grapevine has been a key strategy for developing durable disease-resistant varieties (Xiao et al., 2023). Notably, two members of the TIR-NB-LRR gene family from *Muscadinia rotundifolia* (a close related species of *V. vinifera*) have been identified as major *R* genes, conferring resistance to several important grapevine pathogens (Feechan et al., 2013). Interestingly, it has also been frequently observed that the loss of function of *S* genes can lead to broad-spectrum resistance, making them attractive targets for breeding programs (Zaidi et al., 2018).

Another promising strategy involves triggering grapevine immunity by eliciting plant defense responses with bioproducts (Adrian et al., 2024). Plant innate immunity relies on two well-characterized defense signaling pathways (Boller and Felix 2009). The first one is microbe-associated molecular pattern (MAMP)-triggered immunity (MTI), also referred to as pattern-triggered immunity (PTI). The second is the effector-triggered immunity (ETI) which is activated upon recognition of pathogen-derived effectors by plant resistance proteins (Böhm et al., 2014; Zipfel 2014). MTI represents the initial layer of plant immune defense, involving the recognition of conserved MAMPs released by invading microbes. These molecular patterns are detected by cell-surface pattern recognition receptors (PRRs), which initiate intracellular signal transduction cascades. This signaling typically involves phosphorylation events mediated by mitogen-activated protein kinases (MAPKs), ultimately leading to the activation of transcription factors and the expression of defense-related genes. Depending on the nature and concentration of the MAMPs, various defense responses can be triggered, including callose deposition, production of phytoalexins (such as stilbenes in grapevine), and synthesis of pathogenesis-related (PR) proteins (Ahuja et al., 2012; Zhou and Zhang, 2020).

Plant PRRs are structurally diverse and encompass several protein families, classified based on the nature of their extracellular domains (Ngou et al., 2022). Among these, the Lysin motif receptor-like kinases (LysM-RLKs) and Lysin motif receptor-like proteins (LysM-RLPs) are well-characterized for their ability to perceive various polysaccharides, including chitooligosaccharides (COS), lipochitooligosaccharides (LCOs), and peptidoglycan (PGN). In all plant species, the LysM-RLK family is subdivided into two main groups: LYK receptors, which possess an active kinase domain capable of transducing signals into the cell, and LYR receptors, which contain an inactive kinase domain and are presumed to act as co-receptors. In contrast, LysM-RLPs (also referred to as LYMs) lack a kinase domain altogether and are thought to function in ligand recognition without direct signal transduction (Buendia et al., 2018).

Chitin, a major structural component of fungal cell walls, is perceived by a complex of LysM-RLKs and LysM-RLPs at the plasma membrane, which together mediate signal transduction. In *Arabidopsis thaliana*, the LYR-type receptor AtLYK5, known for its high affinity for chitin oligomers, forms heterodimers with another LYR receptor, AtLYK4 (Wan et al., 2012; Cao et al., 2014; Xue et al., 2019). Despite their ability to bind chitin, the AtLYK4/5 complex cannot initiate intracellular signaling due to their inactive kinase domains. Instead, signal transduction relies on the recruitment of the LYK-type receptor AtCERK1, which possesses an active kinase domain but has relatively low affinity for chitin oligomers (Miya et al., 2007; Cao et al., 2014). In rice (*Oryza sativa*), the LYK-type receptor OsCERK1 plays a dual role in both immune and symbiotic signaling pathways (Zhang et al., 2021). For immune responses, OsCERK1 interacts with LysM-RLPs such as OsCEBiP, OsLYP4, and OsLYP6 to activate defense signaling upon chitin recognition (Shimizu et al., 2010; Ao et al., 2014). In contrast, during the establishment of arbuscular mycorrhizal (AM) symbiosis, OsCERK1 forms a complex with the LYR-type receptor OsMYR1 to mediate symbiotic signaling (He et al., 2019). These studies collectively highlight the central role of OsCERK1 in signal transduction, functioning across distinct biological contexts. The specificity of the signaling pathway depends on the nature of the ligand and the high-affinity interaction with its corresponding receptor.

Phylogenetic analyses have revealed well-conserved clades within the LysM receptor family, reflecting their functional specialization (Buendia et al., 2018). For example, the LYRIII C clade includes AtLYK5 and MtLYR4, which share a conserved function in the perception of chitin oligomers and play a role in MTI activation in *A. thaliana* and *Medicago truncatula*, respectively (Cao et al., 2014; Feng et al., 2019). Similarly, the LYRI A clade, which comprises OsMYR1 and SlLYK10, is functionally conserved in the perception of lipochitooligosaccharides (LCOs) and is involved in the establishment of AM symbiosis in rice and tomato, respectively (Buendia et al., 2016; Girardin et al., 2019; He et al., 2019; Cullimore et al., 2023). In grapevine, recent studies support a similar model in which VvLYK1-1 and VvLYK1-2 (orthologs of AtCERK1), along with VvLYK5-1 (ortholog of AtLYK5), mediate chitin signaling. These LysM-RLKs have been shown to participate in chitin perception and restore MAPK activation, defense gene expression, and pathogen resistance in *A. thaliana* mutants lacking functional chitin receptors (Brulé et al., 2019; Roudaire et al., 2023). Moreover, the formation of a receptor complex between VvLYK1-1 and VvLYK5-1 was found to be dependent on the presence of chitin oligomers. Interestingly, in grapevine, 16 LysM-RLK members have been identified and classified into ten of the eleven phylogenetic clades defined by Buendia et al. (2018), suggesting a broad diversification and potential functional specialization of this receptor family in this species.

In the present study, we investigated the role of VvLYK6, a member of the *V. vinifera* LYRI B LysM-RLK clade, which remains poorly characterized. Orthologs of VvLYK6 in various monocot and dicot species, including tomato, have been implicated in the establishment of AM symbiosis (Li et al., 2022; Ding et al., 2025). Interestingly, Brulé et al. (2019) reported that VvLYK6 was the most highly expressed LysM-RLK during *B. cinerea* infection in susceptible mature grape berries. Using functional genomics approaches, we demonstrated that VvLYK6 suppresses chitin-triggered immune responses and increases plant susceptibility to three distinct fungal pathogens. Our findings suggest that VvLYK6 acts as a negative regulator of plant immunity, contributing to the heightened susceptibility to grey mold. Therefore, *VvLYK6* may be considered as a novel *S* gene encoding a negative regulator of *V. vinifera* immune responses.

## Materials and methods

### Plant materials and growth conditions

*A. thaliana* wild-type (WT) Columbia (Col-0) ecotype, *Atcerk1* mutant (GABI-Kat_096F09, allele *Atcerk1-2*; Gimenez-Ibanez et al., 2009) or transgenic lines Col-0-*p35S::VvLYK6*, Col-0- *p35S::VvLYK6-GFP*, and *Atcerk1*-*p35S::VvLYK6* were grown under a 10/14-h day/night cycle at 20/18°C. Transgenic lines were obtained by floral-dip transformation (Clough and Bent, 1998) of the Col-0 ecotype and *Atcerk1* mutant lines with coding sequence of *VvLYK6* (Vitvi05g00623) amplified from complementary DNA (cDNA) of the susceptible *Vitis vinifera* cv. Marselan leaves and cloned into the pFAST_R02 and pFAST_R05 (Shimada et al., 2010). Transgenic seeds were selected with glufosinate (50 mg/L) and kanamycin (50 mg/L) selection for pFAST_R02 and pFAST_R05, respectively, and subsequently genotyped (Figure S1). All experiments on *A. thaliana* were performed on the third selected homozygous lines.

*V. vinifera* cv. Marselan cells were cultivated in Nitsch-Nitsch (NN) medium as detailed in Roudaire et al. (2023). For experimentations, seven-day-old cultures were diluted twice with new medium. Grapevine cells were transformed by co-cultivation with *Agrobacterium tumefaciens* culture with an absorbance of 0.3 during 2 days. Then, cells were transferred in NN medium with agar (0.7 %) supplemented with 250 mg/L cefotaxime and 50 mg/L kanamycin for selection. After 1 month, two independent lines expressing *35S-VvLYK6-GFP* were selected.

### Analysis of VvLYK6 gene expression in developmental and pathogen-infected grapevine tissues

Microarray and data analysis were performed as described in Kelloniemi et al. (2015) from grapevine (*V. vinifera cv. Marselan)* leaves and berries infected with *B. cinerea* and leaves infected by *P. viticola*. *VvLYK6* expression profile of *V. vinifera* cv. Garganega berries infected with or without *B. cinerea* was obtained from supplemental data of Lovato et al. (2019). Transcriptomic data highlighting differential *VvLYK6* expression profile between pre-egression and egression state in *V. vinifera* cv. Pinot Noir infected with *B. cinerea* were obtained from Haile et al. (2020).

### Phylogenetic analysis of the VvLYKs

Protein sequences for LysM-RLK family were retrieved using BLAST searches, in which A. thaliana family members were used as query sequences against protein sequences from plant species detailed in Table S1. A 1000 bootstrapped phylogenetic tree gathering 96 protein sequences (Table S1) have been constructed with the Maximum Likelihood method and JTT model (Jones et al., 1992). Alignment and phylogenetic analysis were performed with MEGA X software (Kumar et al., 2018).

### Pathogen assays

The *B. cinerea* inoculum was produced by growing the strain BMM on solid medium (V8/2 tomato juice, 1 % agar) in the dark to promote sporulation (Zimmerli et al., 2001). Conidia were isolated and concentrated in water to infect *A. thaliana* leaves with a final concentration of 5x10^4^ conidia/mL diluted in potato dextrose broth medium (PDB, 0.6%). Four-week-old Arabidopsis leaves were cut and placed in survival conditions during *B. cinerea* infection. The *A. brassicicola* inoculum strain MIAE01824 originated from Agroecology unit collection (UMR1347, Dijon, France) was grown 20 days on solid potato dextrose agar medium (PDA, 19 g.L^-1^) supplemented with sucrose (20 g.L^-1^) and CaCO_3_ (30 g.L^-1^) at 20°C in the dark. Four-week-old Arabidopsis leaves were cut and placed in survival conditions during *A. brassicicola* infection. Lesion surfaces caused by these two necrotrophic fungi were measured with imageJ software (https://imagej.net/ij/).

*Erysiphe necator* assays were performed as described by Roudaire et al. (2023). Fungal structures were visualized using a Leica DMA light microscope (magnification x400). For each treatment, 100 germinated spores were evaluated. The percentage of successful epidermal cell penetration was determined based on the presence of a haustorium within the cell or the emergence of secondary hyphae from the appressorium, as previously described by Roudaire et al. (2023). Three biological independent experiments were performed.

### Confocal microscopy

Confocal microscopy was performed using a Leica TCS SP8 multiphoton microscope with a 40X oil-immersion objective. Marselan suspension cells and leaf segments of *A. thaliana* lines expressing *VvLYK6* tagged with GFP were mounted directly or in ultrapure water between slide and coverslips, respectively and observed. For plasma membrane staining, samples were incubated in 10 µM FM4-64 during 10 min prior to observation. Fluorescent markers were excited with the 488 nm laser. GFP and FM4-64 emissions were bandpass filtered at 500-525 nm and between 616 - 694 nm, respectively. Pictures were then treated with LAS X software.

### Elicitors and plant treatment

In this study, we used as elicitors a purified hexamer (DP6) of chitin (GLU436, Elicityl, Crolles, France at a concentration of 0.05 g/L) and flg22 peptide from *X. campestris* pv. campestris strain 305 (QRLSSGLRINSAKDDAAGLAIS) at 10^-6^ or 10^-8^ M, depending on the bioassay (Trdá et al., 2014). For *A. thaliana* treatments, four-weeks old plant leaves were pre-infiltrated with ultrapure water and placed on 6-wells plate containing ultrapure water during 4 h. Then, ultrapure water was replaced by either ultrapure water (control) or elicitors in contact with leaves during 10 min for MAPKs activation and one hour for RT-qPCR experiments before freezing samples in liquid nitrogen.

Grapevine cell suspensions were equilibrated at 0.1 g/mL and shacked under the same conditions as for their culture. Treatment was done in a volume of 20 mL with a final concentration of 0.05 g/L chitin DP6, cells were harvested in liquid nitrogen at 10 min post-treatment for MAPKs activation and 1hour post-treatment for RT-qPCR analysis.

### MAPKs phosphorylation

For MAPK phosphorylation, treated *A. thaliana* leaves or grapevine cells were crushed in liquid nitrogen and total proteins were extracted with a solution containing 50mM HEPES (pH 7.5), 5mM EGTA (pH 8.1), 5 mM EDTA, 1mM Na_3_VO_4_, 50 mM β-glycerophosphate, 10 mM NaF, 1 mM de phenylmethanesulfonyl fluoride, 5 mM dithiothreitol and complete EDTA-free protease inhibitor cocktail (Roche). Phosphorylation of MAPKs was detected by immunoblotting of 25µg total proteins using an anti-p42/44-phospho-ERK antibody (Cell Signaling Technology). Antibody was then revealed using ECL prime (Cytiva) western blotting detection reagent with an Amersham ImageQuant800 (Cytiva). Quantification of MAPKs intensity was normalized by total proteins stained by Ponceau red with the ImageQuant software.

### Real-time quantitative PCR (RT-qPCR) analysis

For RT-qPCR experiments, treated leaves or cells were crushed in liquid nitrogen and total RNA was extracted with the SV Total RNA Isolation System with DNAse treatment (Promega). First-strand cDNAs were synthesized with the High-Capacity cDNA Reverse Transcription kit (Applied Biosystems). RT-qPCR was performed in a ViiA™ 7 Real-Time PCR system (Applied Biosystems) with 10 ng cDNA and 1X GoTaq® qPCR Master Mix (Promega). Relative gene expression was assessed taking into consideration the efficiency (E) of each reaction calculated by the LinRegPCR quantitative PCR data analysis program (Ruijter et al., 2009). Gene expression values were normalized with two housekeeping genes corresponding to *AtRHIP1* (At4g26410) and *AtPTB1* (At3g01150) for RT-qPCR experiments on *A. thaliana* and *VvRPL18B* (Vitvi05g00033) and *VvVPS54* (Vitvi10g01135) for RT-qPCR assays on grapevine cells (*Vitis vinifera* cv. Marselan). All primers used are listed in Table S2.

### Callose deposition analysis

Four-week-old Arabidopsis were sprayed on both sides with either water (as control) or chitin DP6 (0.05 g/L). Four days post-treatment, two leaves of three independent plants per condition were sampled and fixed in absolute ethanol during one night. The aniline blue staining was performed as detailed in Roudaire et al. (2023). Leaves were then observed on adaxial side by epifluorescence microscopy under λex = 340-380 nm and λem = 425 nm, magnification x100, Leica DMRB. Callose deposits were acquired for each condition using the Nis Elements BR software (Nikon) with the DS-5Mc-U1 digital photomicrographic camera (Nikon). Image analysis was then performed on the Fiji application as described in Mason et al. (2020) with Trainable Weka Segmentation plugin.

### Transient expression for FRET-*FLIM analysis*

The transient expression in *Nicotiana benthamiana* leaves were previously described in Girardin et al. (2019) using Agrobacteria containing a cassette overexpressing either VvLYK1-1^G328E^ (Brulé et al., 2019) fused with a C-terminal Cyan Fluorescent Protein (CFP) alone or with VvLYK6 or VvLYK5-1 (Roudaire et al., 2023) tagged with a C-terminal Yellow Fluorescent Protein (YFP) (Karimi et al., 2005). The kinase-dead mutant version of VvLYK1-1 (VvLYK1-1^G328E^) was generated using the Quik Change Site-Directed Mutagenesis protocol (Liu and Naismith, 2008). FLIM was performed on an inverted LEICA DMi8 microscope equipped with a TCSPC system from PicoQuant. The excitation of the CFP donor at 440 nm was carried out by a picosecond pulsed diode laser at a repetition rate of 40 MHz, through an oil immersion objective (63×, N.A. 1.4). The emitted light was detected by a Leica HyD detector in the 450-500 nm emission range. Images were acquired with acquisition photons of up to 1500 per pixel. Data were acquired in 3 independent experiments and pooled. From the fluorescence intensity images, the decay curves of CFP were calculated per pixel and fitted (by Poissonian maximum likelihood estimation) with either a double- or tri-exponential decay model using the SymphoTime 64 software (PicoQuant, Germany). The double-exponential model function was applied for donor samples with only CFP present. The tri-exponential model function was used for samples containing CFP and YFP.

### Co-immunopurification with VvLYK1-1

For co-immunopurification, the sequence coding VvLYK1-1^G328E^ was fused to the sequence coding a triple hemaglutin (HA) tag. VvLYK1-1^G328E^-HA and VvLYK6-YFP or VvLYK5-1-YFP were expressed into *N. benthamiana*. Leaf material was collected 3 days after infiltration and co-immunopurification was performed as described in Ding et al. (2024), except that anti-HA beads (pierce Anti-HA magnetic beads, 88836) were used for purification and anti-HA-HRP antibodies (Roche, Anti-HA-Peroxidase High affinity from rat IgG1, 12013819001) were used for Western-blotting. Chitin DP6 (CO6) at 0.1 g/L was infiltrated into leaves before harvest and added in the IP buffer for protein solubilisation and wash of the beads.

### Microsomal fraction preparation and binding assay

Preparation of microsomal fraction, and binding assays were previously described in Girardin et al., 2019. CO5-biotin and CO7-biotin were synthesized as detailed in Cullimore et al., 2023. Briefly, binding assays with CO5-biotin and CO7-biotin were conducted on microsomal fractions in binding buffer described in Girardin et al. (2019) for 1h at 4 °C. For western-blotting, proteins were separated by SDS-PAGE using homemade 6 % polyacrylamide gels and transferred onto a nitrocellulose membrane with a transblot system (Bio-rad). The membranes were blocked for 1h and incubated for 1h with the following antibodies: αGFP (11814460001, Roche, 1:3000), and then goat αMouse-HRP (1706516, Bio-Rad, 1:10000) or streptavidin-HRP (S911, Invitrogen, 1:3000). The chemiluminescent signal from HRP was observed using Chemidoc system (Bio-Rad).

## Results

### Analysis of VvLYK6 expression suggests an important role during B. cinerea infection

To investigate the role of VvLYK6 in grapevine immunity, we examined its expression profile using our previously published transcriptomic dataset (Kelloniemi et al., 2015). Our analysis revealed that *VvLYK6* is upregulated in susceptible *V. vinifera* leaves at 24 and 48h post-infection (hpi) with *B. cinerea*, whereas its expression remains unchanged following infection with *P. viticola* (Figure 1A). Given that VvLYK6 belongs to the LysM-RLK family known to perceive COS, these results are consistent with the presence of chitin in the fungal cell wall of *B. cinerea*, which is nearly absent in that of the oomycete *P. viticola*. However, *VvLYK6* expression was not significantly induced in grapevine cells treated with chitin hexamers (Figure S2A). We also observed that *VvLYK6* expression is induced after 48h of *B. cinerea* infection on grapevine fruits (Figure 1B) and particularly in infected mature berries (Figure 1C) which are known to be very susceptible to grey mold (Kelloniemi et al., 2015). To corroborate these results and confirm the upregulation of *VvLYK6* during *B. cinerea* infection, we analyzed other publicly available transcriptomic data (Figure S2B-C). In a first study on mature berries of the susceptible *V. vinifera* cv. Garganega inoculated or not with *B. cinerea*, *VvLYK6* expression was significantly increased in infected berries compared to uninfected ones twelve days post-inoculation (Lovato et al., 2019, Figure S2B). In a second study comparing ripe berries of the susceptible *V. vinifera* cv. Pinot Noir with and without visible symptoms of *B. cinerea* infection (referred to as egression and pre-egression, respectively), the expression of *VvLYK6* was significantly upregulated only in berries displaying visible symptoms (Haile et al., 2020, Figure S2C). All together, these results showed a correlation between *VvLYK6* upregulation in mature grapevine berries and their high susceptibility to *B. cinerea*.

**Figure 1.**
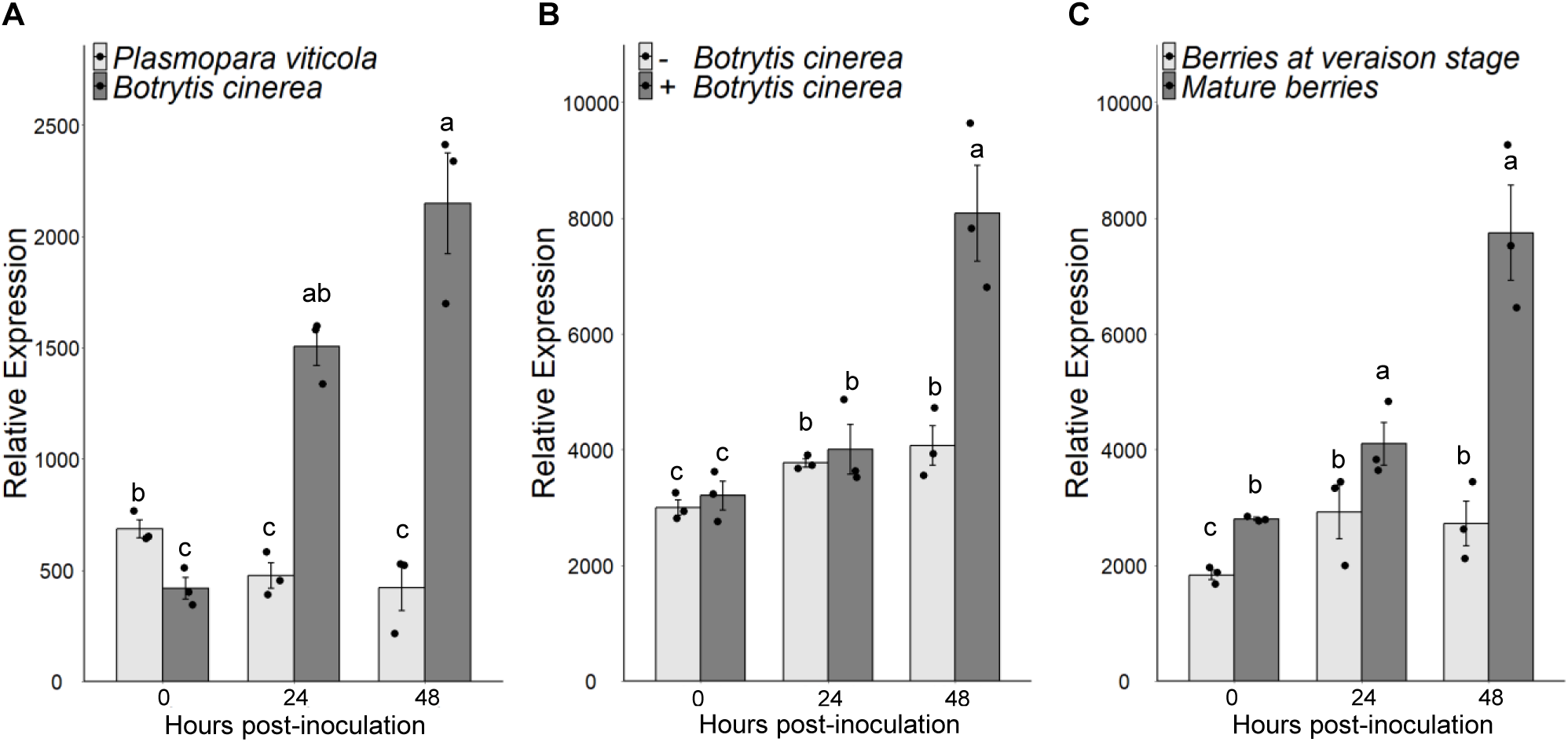
*VvLYK6* is induced during *Botrytis cinerea* infection on grapevine leaves and mature berries. **(A)** *VvLYK6* expression profiles analyzed at 0, 24 and 48 h post-inoculation on leaves infected either by *Botrytis cinerea* or *Plasmopara viticola*. **(B)** *VvLYK6* expression profiles analyzed at 0, 24 and 48 h post-inoculation on ripe berries infected or not with *B. cinerea*. **(C)** Comparison of *VvLYK6* expression on grapevine berries infected with *B. cinerea* at two stages of grapevine development corresponding to berries at veraison stage and ripe berries, respectively. For each bar plot, bars represent the mean ± SEM on three biologically independent experiments and statistically significant groups are categorized by different letters (Kruskal-Wallis/BH post-test; P<0.05). All transcriptomic data were generated from Kelloniemi et al. (2015).

### VvLYK6 belongs to the LYRI B clade with conserved domain of LysM-receptor like kinase

To investigate the function of VvLYK6, we first conducted a phylogenetic analysis involving several plant species known to possess well-characterized LysM-RLKs associated with immunity and/or symbiosis. LysM-RLK protein sequences were retrieved from *A. thaliana*, *Hordeum vulgare, Oryza sativa*, *Medicago truncatula*, *Musa acuminata*, *Prunus persica*, *Solanum lycopersicum* and *V. vinifera* (Supplementary Table S1). A total of 96 protein sequences were selected to construct a maximum-likelihood phylogenetic tree (Figure 2A). The resulting tree displays the two well-known sub-groups (LYKs and LYRs) further divided in eleven clades as described by Buendia et al. (2018). The 16 members of the LysM-RLK family in *V. vinifera* are distributed across all clades except for the LYKIV clade, which is not conserved among dicots (Figure 2A). Our study focused on VvLYK6 (indicated by a black arrow), which belongs to the LYRI B clade, a group whose function has never yet been characterized in *V. vinifera*.

**Figure 2.**
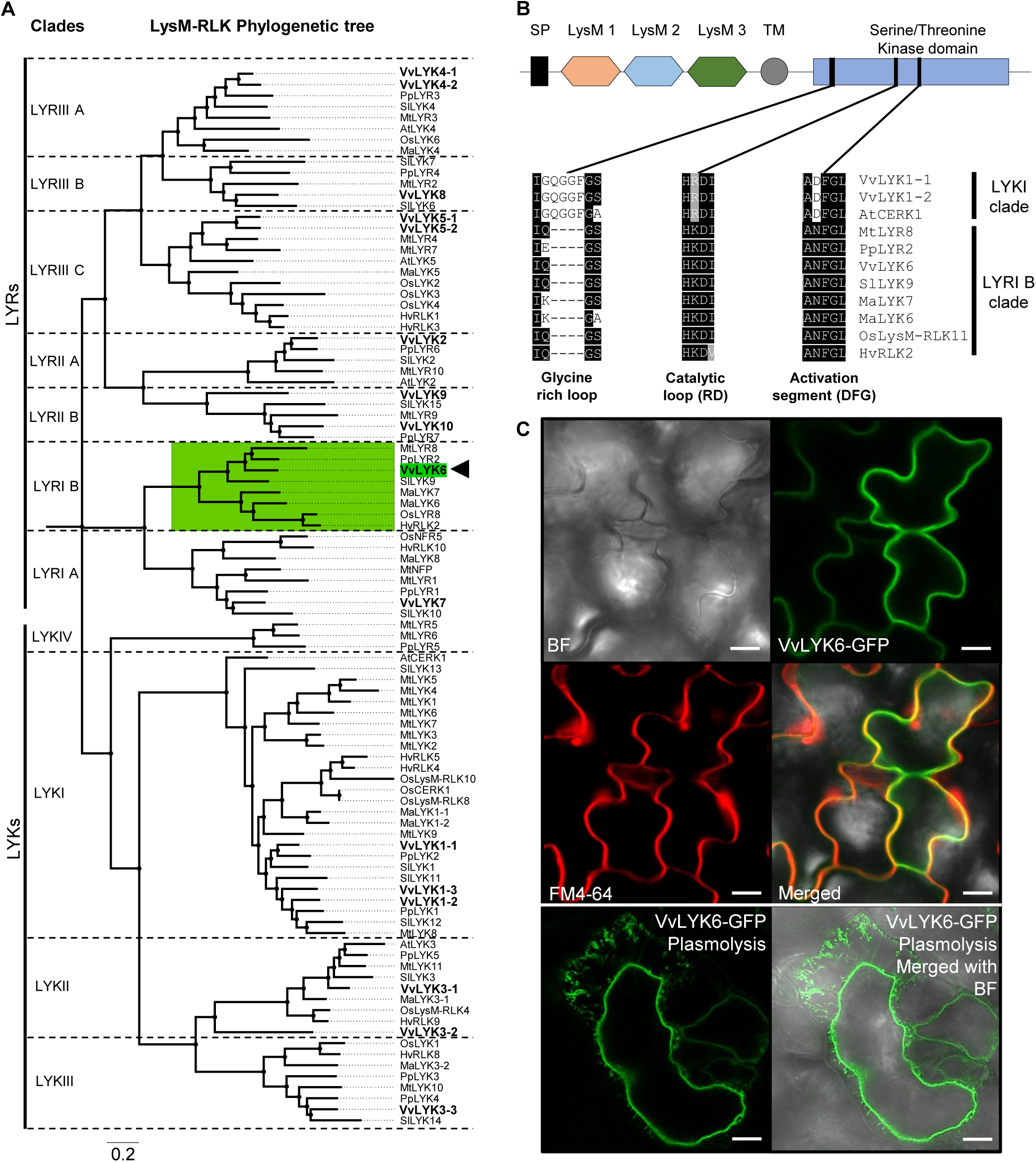
*VvLYK6* belongs to the LYRI B phylogenetic clade of the LysM receptor-like kinases family. **(A)** A 1000 bootstrapped Maximum-likelihood phylogenetic tree (Jones-Taylor-Thornton (JTT) model) with LysM-RLKs from 8 plant species was constructed with MEGA X (Kumar *et al*., 2018). Phylogenetic tree was subdivided by clades according to the phylogenetic analysis from Buendia *et al*. (2018). LYKs correspond to LysM-RLKs with an active kinase domain and LYRs correspond to LYK-related LysM-RLKs without kinase activity. Protein sequences are listed in supplemental Table S1. The 16 LysM-RLKs found in grapevine genome (VvLYKs) are in bold. VvLYK6 is highlighted in green and indicated by a black arrow inside the green-surrounded LYRI B clade. **(B)** Schematic representation of putative VvLYK6 domains. Black square: Signal peptide, hexagon: LysM domain, LysM1 (orange), LysM2 (blue), LysM3 (green), grey circle: transmembrane domain and blue square: kinase domain. Below schematic VvLYK6 structure are shown alignments of motifs involved in kinase activity (Glycine rich-loop, Catalytic loop (RD) and activation segment (DFG motif). Protein domain sequences were aligned with Clustal performed on MEGAX (Kumar *et al*., 2018). Black and grey shading representing conservation of amino acids. The alignment includes VvLYK1-1, VvLYK1-2, AtCERK1 belonging to the LYKI clade and MtLYR8, PpLYR2, VvLYK6, SlLYK9, MaLYK7, MaLYK6, OsLysM-RLK11/OsLYR8 and HvRLK2 belonging to the LYRI B clade. **(C)** Subcellular localization of VvLYK6 in *A. thaliana* leaves. GFP-tagged VvLYK6 in leaf segments of *A. thaliana* co-localizes with the plasma membrane marker probe (FM4-64). Plasmolysis of plant cells expressing VvLYK6 tagged with GFP reveals that the GFP signal does not localize to the cell wall. Scale bars represent 20 µm. Similar localization was observed in three independent lines. BF = Brightfield.

We began by analyzing the putative structure of VvLYK6, confirming the presence of all conserved domains typical of LysM-RLKs: a signal peptide, three extracellular LysM domains, a transmembrane domain, and a cytoplasmic putative serine/threonine kinase domain (Figure 2B, Table S3). Regarding the intracellular kinase domain, most of the amino acids of the kinase domain present in LYKI members (e.g. AtCERK1, VvLYK1-1, VvLYK1-2) were conserved in LYRI B orthologs, including VvLYK6 (Figure 2B). However, deletions observed in the glycine-rich loop, along with the lack of conservation in the catalytic loop and activation segment (both essential for kinase activity) suggest that the kinase domain of VvLYK6, as well as other members of the LYRI B may be catalytically inactive (Hanks et al., 1988; Klaus-Heisen et al., 2011). Specifically, while the glycine-rich loop typically contains three to four conserved glycine residues in most LYK subgroup members with kinase activity, only a single glycine residue is present in LYRI B members. Furthermore, other key motifs associated with kinase function, such as the catalytic loop (RD) and the activation segment (DFG), which are well conserved in active LYK kinases, are absent in the LYRI B clade (Figure 2B).

To determine the subcellular localization of VvLYK6, we expressed a C-terminal GFP-tagged version of the protein in *A. thaliana* and *V. vinifera*. Transgenic lines exhibited green fluorescence from VvLYK6-GFP which co-localized with the red fluorescence of the plasma membrane marker FM4-64 (Brandizzi et al., 2004) in both plant species (Figure 2C, Figure S3). Plasmolysis experiments further confirmed that VvLYK6 followed the plasma membrane during cell shrinkage (Figure 2C) supporting its predicted localization at the plasma membrane.

### Constitutive expression of VvLYK6 in A. thaliana increases its susceptibility to B. cinerea and other adapted or non-adapted fungal pathogens

To investigate the role of VvLYK6 during *B. cinerea* infection, and taking advantage of the fact that *VvLYK6* has no ortholog in *A. thaliana*, we constitutively expressed its gene from the susceptible *V. vinifera* cv. Marselan under the control of the *CaMV35S* promoter (*p35S*) in the wild-type (WT) ecotype Col-0 of *A. thaliana*. Three independent lines exhibiting high levels of *VvLYK6* transcripts were selected (Figure S4A). To assess the impact of *VvLYK6* expression during fungal infections, leaves were inoculated with *B. cinerea* and lesion areas were quantified three days post-inoculation (dpi). As shown in Figure 3A and Figure S4B, all three transgenic lines displayed a significant increase in susceptibility to *B. cinerea*, with lesion areas approximately 50% larger than those observed in WT plants.

**Figure 3.**
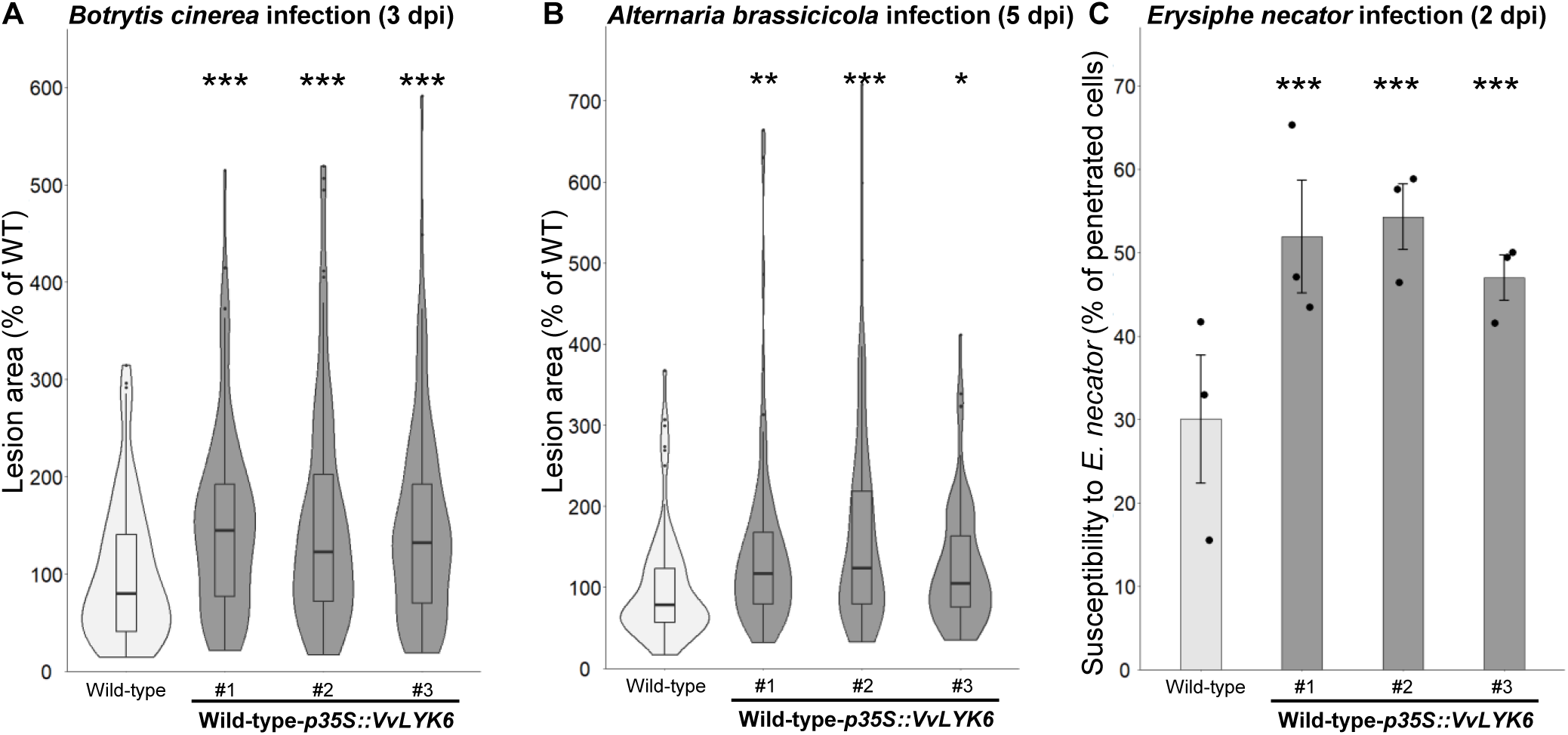
Constitutive expression of *VvLYK6* in *A. thaliana* increases its susceptibility to different fungal pathogens. **(A)** Disease lesions percentage induced by *B. cinerea* on Wild-type (WT) and the three independent transgenic lines expressing *VvLYK6* measured 3 days post inoculation, lesion area on WT plants are referred as 100%. Violin plot with included boxplot illustrates the distribution of lesions percentage of three independent experiments. For each biological replicate, 32 lesions were scored from 8 plants per line, combining data from 3 biological replicates. Each transgenic line was statistically compared to the WT with the non-parametric Wilcoxon test (***, P<0.001). **(B)** Lesions percentage provoked by the necrotrophic fungus *A. brassicicola* on the WT and the three independent transgenic lines constitutively expressing *VvLYK6,* measured five days post inoculation. Violin plot with included boxplot represents distribution of lesions percentage of five independent experiments. For each biological replicate, 18 lesions were scored from four plants per line, combining data from 4 biological replicates. Each transgenic line was statistically compared to the WT with the non-parametric Wilcoxon test (***, P<0.001; **, P<0.01; *, P<0.05). **(C)** Penetration of the non-adapted powdery mildew *E. necator* on the WT and the three independent transgenic lines constitutively expressing *VvLYK6* measured 2 days post inoculation. Bars represent the mean percentage of epidermal penetrated cells from three biologically independent experiments each with more than one hundred germinated conidia counted ± SEM. Each transgenic line was statistically compared to the WT with a pairwise comparison of proportions test (***, P<0.001).

We next examined whether VvLYK6 expression influenced susceptibility to other fungal pathogens. First, we tested the necrotrophic fungus *Alternaria brassicicola*, which is adapted to Brassicaceae. At 5 dpi, the three transgenic lines showed significantly larger lesion areas, ranging from 26% to 67%, compared to WT plants (Figure 3B, Figure S4C).

Finally, we evaluated the response to the non-adapted biotrophic fungus *Erysiphe necator* (grapevine powdery mildew). The Arabidopsis transgenic lines expressing *VvLYK6* exhibited a significantly higher rate of epidermal cell penetration compared to WT plants, with approximately 50% of cells penetrated in transgenic lines versus 30% in controls (Figure 3C).

All these results suggest that VvLYK6 may interfere with plant immune responses, thereby facilitating fungal colonization and spread.

### Constitutive expression of VvLYK6 inhibits chitin-triggered immune responses in A. thaliana

To further elucidate the function of VvLYK6, we investigated early defense responses triggered by chitin by analyzing MAPK phosphorylation levels. As previously reported (Roudaire et al., 2023), chitin oligomers induce transient phosphorylation of two mitogen-activated protein kinases (MAPKs) of 43 and 47 kDa in *A. thaliana* WT plants (Figure 4A), corresponding to MPK3 and MPK6, respectively (Claverie et al., 2018). Interestingly, phosphorylation of both MAPKs, measured 10 min after treatment with chitin hexamers (DP6), was significantly reduced, by approximately 40%, in all three independent *VvLYK6*-expressing lines compared to WT plants (Figure 4A–B). To rule out a delayed activation of MAPKs in transgenic lines, phosphorylation levels were also assessed 20 min post-treatment, but no compensatory increase was observed (Figure 4A).

**Figure 4.**
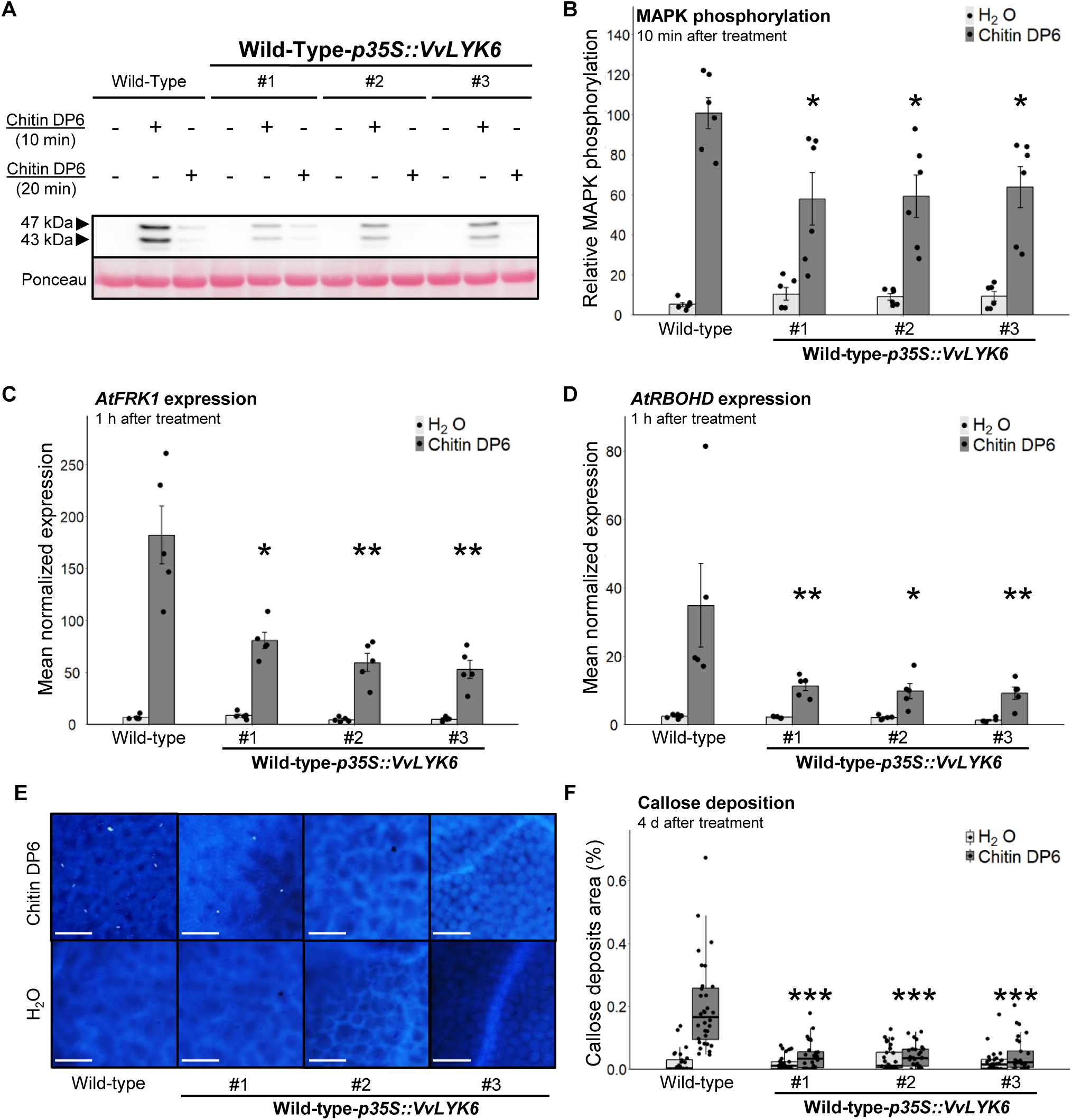
Constitutive expression of VvLYK6 inhibits chitin-triggered immune responses in *A. thaliana*. **(A-B)** MAPKs phosphorylation was detected 10 min or 20 min after H_2_O (-) or chitin DP6 treatment (0.05 g/L) by immunoblotting with an antibody α- pERK1/2 in the Wild-Type (WT) and three independent lines constitutively expressing *VvLYK6*. (a) Representative immunoblotting with α- pERK1/2 and homogeneous loading verified by Ponceau red staining. **(B)** Quantification of the MAPKs phosphorylation at 10 min after treatment, detected by ImageQuant. Bars represent the mean ± SEM on 6 biologically independent experiments. Each *VvLYK6*-expressing line treated with chitin DP6 was statistically compared to the WT with the non-parametric Wilcoxon test (*, P<0.05). **(C-D)** Normalized expression level of *AtFRK1* and *AtRBOHD* measured by qPCR 1h after chitin DP6 (0.05 g/L) or H_2_O treatment. The expression levels of *AtFRK1* and *AtRBOHD* were normalized to those of two housekeeping genes. Bars represent the mean of relative expression ± SEM of 5 biologically independent experiments. Asterisks indicate a statistically significant difference between each transgenic line and the WT treated with chitin DP6, (non-parametric Wilcoxon test; *, P<0.05; **, P<0.01). **(E)** Representative pictures of callose deposition after chitin DP6 (0.05g/L) or H_2_O treatment in WT or the three *VvLYK6*-expressing lines, observed 4 days post-treatment and analyzed by epifluorescence microscopy after aniline blue staining. Callose deposits were quantified with Trainable Weka Segmentation plugin in imageJ. **(F)** Box plot represents distribution of callose deposits percentage on total leaves surface for each genotype and condition on at least 30 pictures per condition from three biologically independent experiments. Asterisks indicate a significant difference between each line expressing *VvLYK6* and WT treated with DP6 (Wilcoxon test, ***; P<0.001).

In addition, we examined the expression of two defense-related genes: *AtFRK1* (flagellin-induced receptor kinase 1) and *AtRBOHD* (respiratory burst oxidase homolog D), both known to be induced by chitin DP6 treatment (Morales et al., 2016; Brulé et al., 2019). Expression of *AtFRK1* was significantly reduced in the three transgenic lines, with transcript levels decreased by 56% to 71% compared to WT (Figure 4C). Similarly, *AtRBOHD* transcript levels were reduced by 67% to 74% in the *VvLYK6*- expressing lines (Figure 4D). To further explore the function of VvLYK6, we analyzed additional immune-related genes involved in hormonal signaling pathways, such as salicylic acid (SA) and jasmonic acid (JA) (Spoel et al., 2003). No significant differences were observed in the expression of the SA- and JA-responsive genes *PR1* and *PDF1.2* between the three independent *VvLYK6*-expressing lines and WT plants following chitin DP6 treatment (Figure S5).

We then examined the impact of VvLYK6 expression on callose deposition, a later-stage defense response. Chitin is well known to induce callose accumulation in *A. thaliana* leaves (Roudaire et al., 2023). Compared to WT plants, all transgenic lines expressing *VvLYK6* exhibited a significantly reduced percentage of callose deposits four days after chitin DP6 treatment (Figure 4E–F). Specifically, WT leaves showed callose deposits covering 0.2% of the total leaf area, whereas the transgenic lines averaged only 0.04%, a level comparable to water-treated controls (Figure 4E–F).

Taken together, these results clearly indicate that VvLYK6 expression in *A. thaliana* suppresses chitin-triggered immune responses.

Although our study primarily focused on the role of VvLYK6 during fungal infection and chitin perception, given its induction during *B. cinerea* infection, we also tested its specificity by analyzing responses to a bacterial MAMP: the flagellin-derived peptide flg22 (Trdá et al., 2014). MAPK phosphorylation was assessed 10 min after flg22 treatment in WT and *VvLYK6*-expressing lines. Immunoblot analysis revealed no significant differences between the transgenic lines and WT (Figure S6A). As flg22 is also known to inhibit plant growth in *A. thaliana* (Vetter et al., 2012), flg22-induced growth inhibition assays were performed. After 12 days of growth on a medium containing 1 µM flg22, no significant differences in growth inhibition were observed between WT and *VvLYK6*-expressing seedlings (Figure S6B).

These findings demonstrate that VvLYK6 expression in *A. thaliana* does not interfere with flg22- triggered immune signaling, but appears to specifically modulate chitin-induced immune responses associated with fungal pathogens.

### VvLYK6 represses chitin-triggered immunity in grapevine (Vitis vinifera)

To validate the findings observed in *A. thaliana*, we investigated the function of VvLYK6 in *V. vinifera*. Two independent grapevine cell lines constitutively expressing VvLYK6 tagged with GFP and exhibiting high transcript and protein levels were selected (Figure S7A–B).

Chitin oligomers are known to elicit phosphorylation of two MAPKs (45 and 49 kDa) in grapevine cells (Brulé et al., 2019). Compared to WT cells, MAPK phosphorylation induced by chitin DP6 was significantly reduced by 25% to 35% in both transgenic lines (*VvLYK6* OE#1 and OE#2) after 10 min of treatment (Figure 5A, Figure S7C). We also quantified the expression of two defense-related genes involved in the stilbene biosynthetic pathway, which contributes to phytoalexin production in grapevine: *VvPAL* (phenylalanine ammonia lyase) and *VvSTS1.2* (stilbene synthase). One hour after chitin DP6 treatment, the constitutive expression of *VvLYK6* significantly impaired the induction of both genes. The typical upregulation of *VvPAL* and *VvSTS1.2* was reduced by more than 50% in the transgenic lines compared to WT cells (Figure 5B–C). On the whole, these results suggest that VvLYK6 acts as a negative regulator of chitin-triggered immune responses in *V. vinifera*, consistent with the observations made in *A. thaliana*.

**Figure 5.**
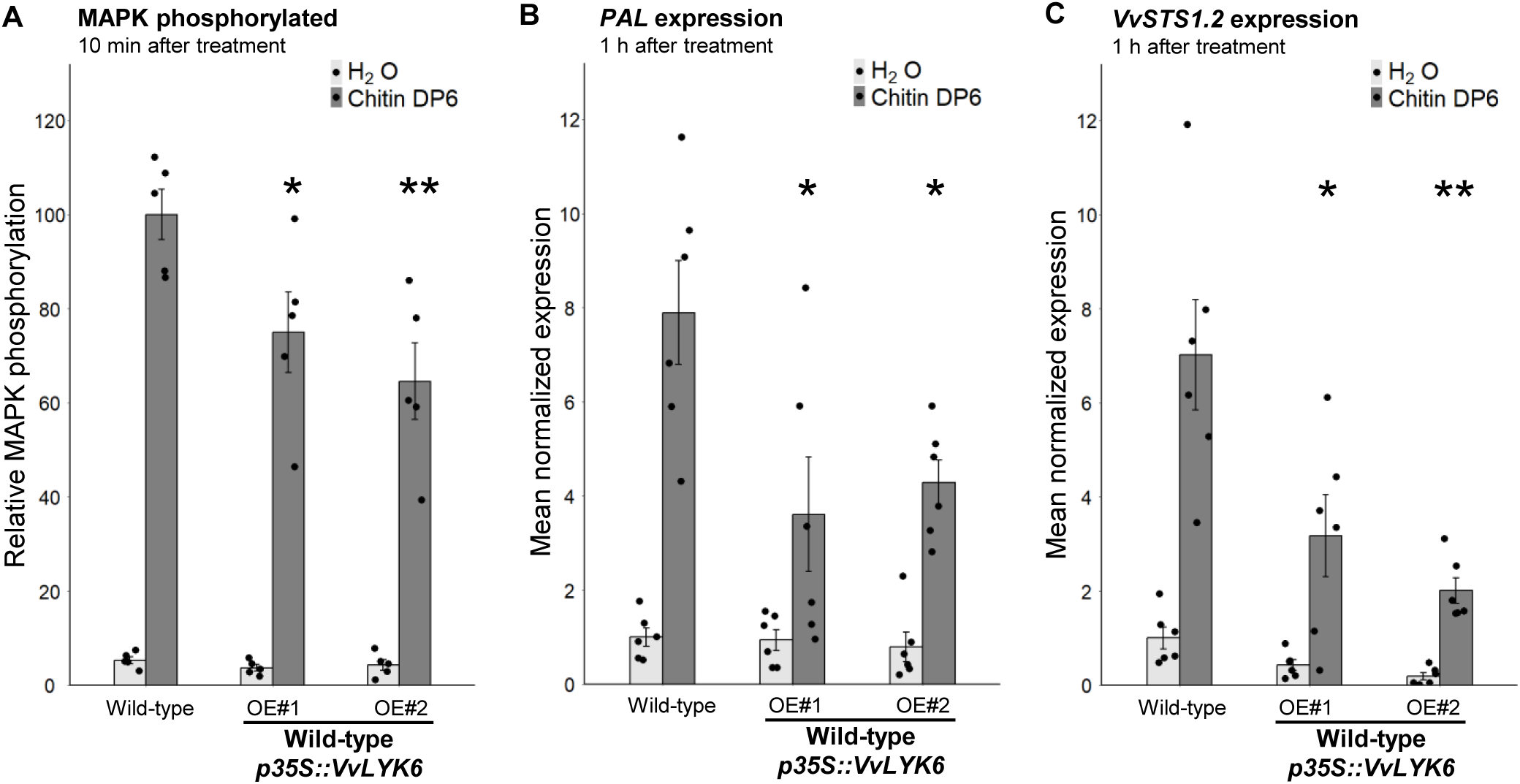
VvLYK6 is also a repressor of chitin-triggered immune responses in *V. vinifera*. **(A)** Quantification of the MAPKs phosphorylation detected 10 min after H_2_O or DP6 treatment (0.05 g/L) by immunoblotting with an antibody α-pERK1/2 in the WT grapevine cell line and two independent lines constitutively expressing VvLYK6-GFP. MAPKs phosphorylation was quantified by ImageQuant. Bars represent the mean ± SEM on five biologically independent experiments. Each *VvLYK6*-expressing cell line was statistically compared to the corresponding WT grapevine cells treated with DP6 using the non-parametric Wilcoxon test (*, P<0.05; **, P <0.01). **(B-C)** Normalized expression level of *VvPAL* **(B)** and V*vSTS1.2* **(C)** measured by qPCR 1h after DP6 (0.05 g/L) or H_2_O treatment. The expression levels of *VvPAL and VvSTS1.2* were normalized to those of two housekeeping genes. Bars represent the mean of relative expression ± SEM of 6 biologically independent experiments. Asterisks indicate a statistically significant difference between each transgenic line and the WT treated with DP6 using a pairwise Wilcoxon test (*, P<0.05; **, P <0.01).

### VvLYK6 forms a receptor complex with VvLYK1-1 in presence of chitin oligomers

During fungal infection or in the presence of chitin, VvLYK1-1 has been identified as a key co-receptor involved in chitin-triggered immune signaling in grapevine (Brulé et al., 2019). Notably, VvLYK1-1 is the only LysM-RLK shown to form a receptor complex with VvLYK5-1 to perceive chitin and activate downstream defense responses (Roudaire et al., 2023).

To explain the repression of microbe-triggered immunity (MTI) observed when *VvLYK6* is overexpressed, we hypothesized a potential interaction between the functional co-receptor VvLYK1-1 and VvLYK6. To test this, we performed FRET-FLIM experiments following transient co-expression of both proteins fused to CFP and YFP, respectively. Since constitutive expression of VvLYK1-1 induces strong cell death in *Nicotiana benthamiana*, we used a kinase-dead version carrying the G328E mutation (VvLYK1-1^G328E^-CFP). As expected, the CFP lifetime of VvLYK1-1^G328E^-CFP expressed alone was unaffected by chitin treatment (Figure 6A). However, co-expression with VvLYK6-YFP followed by chitin DP6 treatment resulted in a significant decrease in CFP lifetime, indicating a ligand-dependent *in vivo* interaction between these two LysM-RLKs (Figure 6A). This interaction was further confirmed by co-immunoprecipitation (Co-IP) assays, which demonstrated that VvLYK6-YFP physically associates with VvLYK1-1^G328E^-HA in the presence of chitin DP6 (Figure 6B). In both experiments, VvLYK5-1 was used as a positive control, as its interaction with VvLYK1-1 in a ligand-dependent manner has been previously described (Figure 6A–B; Roudaire et al., 2023). Interestingly, VvLYK6 and VvLYK1-1 primarily interact in the presence of chitin DP6, suggesting that VvLYK6 may possess affinity for chitin oligomers. Indeed, some LYRI B orthologs have been reported to bind both short- and long-chain chitin oligomers with high affinity (Li et al., 2022; Ding et al., 2025). To evaluate the binding capacity of VvLYK6, we used cross-linkable biotinylated versions of chitin pentamer and heptamer (CO5-biotin and CO7-biotin) to assess its interaction with chitin oligomers. Surprisingly, unlike its orthologs in *Brachypodium distachyon* (BdLYR2) and *Medicago truncatula* (MtLYR8), VvLYK6 did not exhibit high-affinity binding to either CO5 or CO7 (Figure 6C; Li et al., 2022; Ding et al., 2025). This discrepancy may be due to improper folding of VvLYK6 when overexpressed in *Nicotiana benthamiana* leaves, resulting in a non-functional protein. Alternatively, VvLYK6 may require association with VvLYK1-1 to properly bind chitin oligomers and initiate signal transduction. Supporting this last hypothesis, constitutive expression of VvLYK6 in the *Arabidopsis* mutant *Atcerk1* failed to activate MAPK phosphorylation or induce *FRK1* expression following chitin DP6 treatment (Figure S8). These findings suggest that VvLYK6 may not function as a primary chitin receptor but could act as a modulator of immune signaling through interaction with VvLYK1-1.

**Figure 6.**
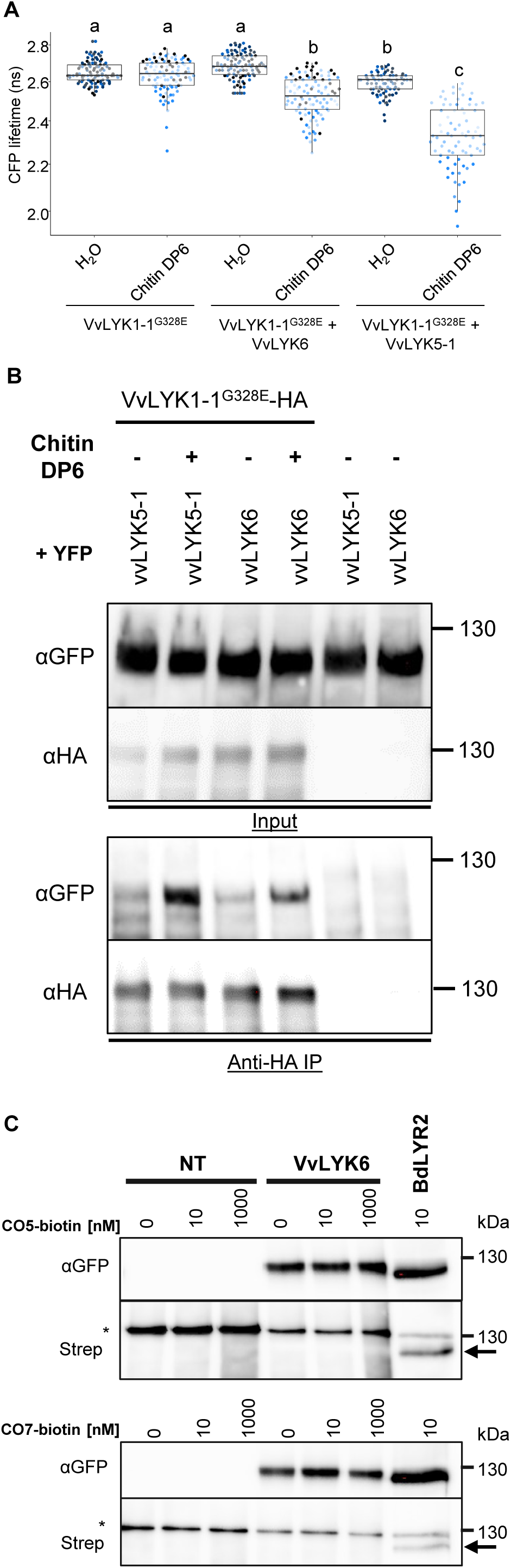
VvLYK6 interacts with VvLYK1-1 in presence of chitin oligomers. **(A)** The dead kinase VvLYK1-1^G328E^-CFP was transiently expressed in *Nicotiana benthamiana* leaves alone or together with VvLYK6-YFP or VvLYK5-1-YFP, a protein known to interacts with VvLYK1-1 in presence of chitin DP6 (Roudaire *et al*., 2023). CFP lifetime (ns) was measured after leaves treatment with either water (H_2_O) or chitin DP6 (0,1 g/L). Boxplots represent the distribution of the data acquired in at least three biologically independent experiments. Different letters indicate a significant difference using a non-parametric Kruskal-Wallis test. **(B)** Co-immunopurification (Co-IP) of VvLYK1-1^G328E^-HA with VvLYK5-1:YFP and VvLYK6:YFP in presence or not of chitin DP6. Extracted proteins from *N. benthamiana* leaves were purified using anti-HA beads (Anti-HA IP). After IP, corresponding proteins were detected with anti-GFP antibodies (αGFP) and anti-HA antibodies (αHA). **(C)** The LysM-RLK VvLYK6 alone is not able to bind CO5 or CO7 with high affinity. VvLYK6- YFP and BdLYR2-YFP were transiently expressed in *N. benthamiana* leaves. Subsequently, microsomal fractions were prepared, and binding assays were conducted using a range of CO5-biotin or CO7-biotin as ligand. Microsomal fractions from leaves expressing BdLYR2-YFP were used as positive control (black arrow) while those from non-transformed (NT) leaves served as negative control. Western blotting was conducted using αGFP and streptavidin-HRP. The quantities of microsomal fractions used correspond to 50 µg of total proteins for VvLYK6-YFP and NT, and 5 µg for BdLYR2-YFP. The asterisk represents an unspecific band (endogenously biotinylated protein) observed at 130 KDa. No binding with CO5-biotin or CO7-biotin were observed with VvLYK6.

## Discussion

*B. cinerea* is among the most devastating fungal pathogens affecting a wide range of plant species. This necrotrophic fungus is particularly known for infecting mature berries of susceptible *V. vinifera* cultivars, leading to significant reductions in yield and must quality during winemaking (Fillinger and Elad, 2016). Numerous transcriptomic studies have highlighted the upregulation of specific genes during berry ripening, a developmental stage highly susceptible to infection (Kelloniemi et al., 2015; Lovato et al., 2019; Haile et al., 2020).

Chitin, a major structural component of fungal cell walls, is recognized as a MAMP. In the context of *B. cinerea* infection, we focused on the well-characterized LysM-RLK family, which plays a key role in the in the perception of chitin oligomers and the activation of immune responses. In susceptible *V. vinifera* varieties, *VvLYK6* has been identified as the most highly expressed LysM-RLK during *B. cinerea* infection in mature berries (Kelloniemi et al., 2015; Brulé et al., 2019), a finding further supported by additional transcriptomic analyses (Lovato et al., 2019; Haile et al., 2020).

Our study provides new insights into the role of *VvLYK6*, a member of the newly defined LYRI B subfamily. Unexpectedly, we found that the constitutive expression of *VvLYK6* in *A. thaliana* significantly increases the plant susceptibility to *B. cinerea* and two other adapted and non-adapted fungal pathogens. More specifically, our results indicate that the inhibitory function of *VvLYK6* is confined to the chitin-triggered defense pathway associated with fungal pathogens.

In *V. vinifera*, the constitutive expression of *VvLYK6* similarly inhibits the chitin-triggered immune pathway. Morevoer, VvLYK6 has been shown to form a heterodimeric complex with VvLYK1-1 in the presence of chitin oligomers, suggesting potential competition between VvLYK6 and VvLYK5-1 for complex formation with VvLYK1-1 (Roudaire et al., 2023). During *B. cinerea* infection in mature berries, the upregulation of *VvLYK6* may lead to a predominance of VvLYK1-1/VvLYK6 complexes, especially since *VvLYK6* expression does not affect *VvLYK5-1* transcript levels (Figure S7D). This could negatively regulate the formation of *VvLYK1-1/VvLYK5-1* complexes, supporting a model in which *VvLYK6* expression modulates chitin-triggered immunity in grapevine (Figure 7). A similar regulatory mechanism has been described in rice within the LysM-RLK family. The rice ortholog of VvLYK1-1, OsCERK1, is known to participate in both immune responses, via interaction with *OsCEBIP*, and AM symbiosis via interaction with OsMYR1. Notably, constitutive expression of *OsMYR1*, a member of the LYRI A clade, significantly reduces disease resistance in rice (Zhang et al., 2021). Analogous regulatory dynamics have also been observed in other RLK families. For instance, the Leucine-Rich Repeat RLK *BIR2* acts as a negative regulator by forming a complex with *BAK1*, thereby preventing *BAK1* from associating with ligand-binding receptors such as *FLS2* in the presence of MAMPs (Halter et al., 2014).

**Figure 7.**
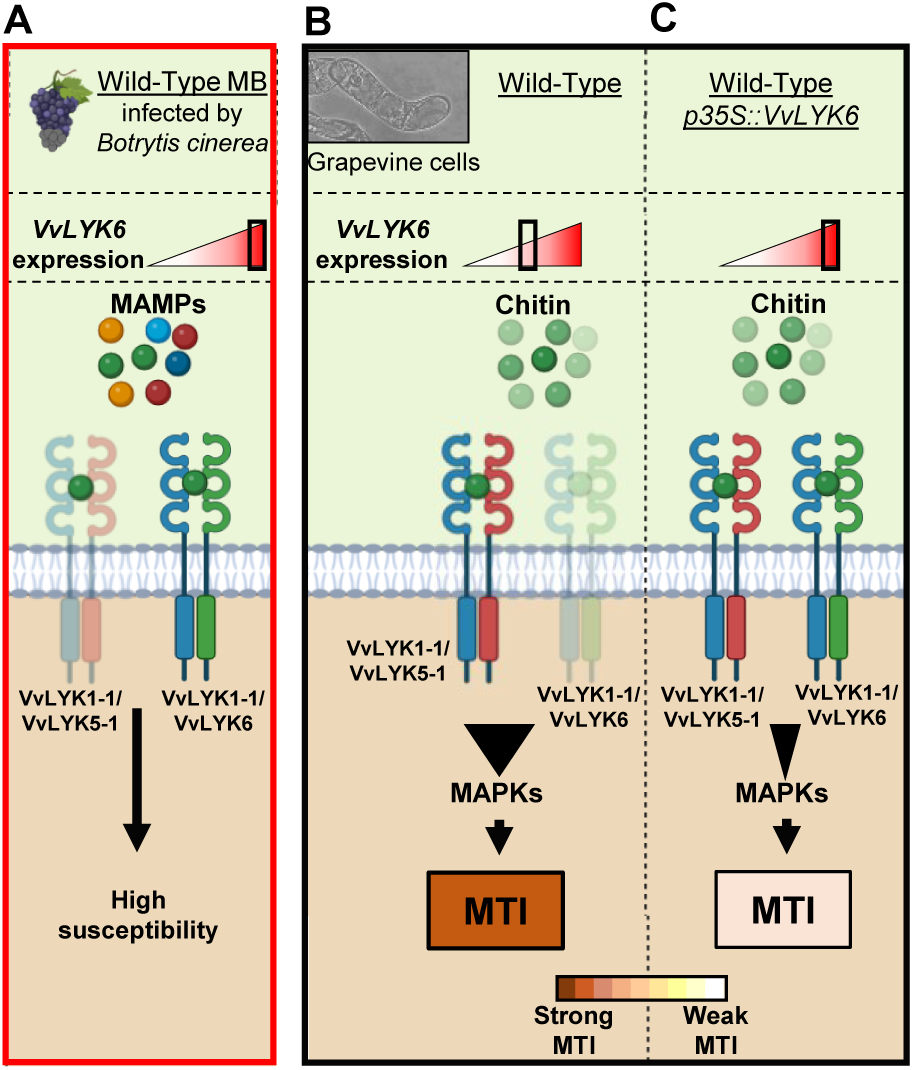
Proposed mode of action of VvLYK6 to modulate plant immune responses. VvLYK6 is involved in grapevine susceptibility during *Botrytis cinerea* infection and repression of chitin oligomer elicitation. **(A)** *VvLYK6* expression is induced during *Botrytis cinerea* infection in mature berries (MB), correlated with the absence of defense gene (*VvSTS1.2* and *VvPAL*) and a high susceptibility to fungal pathogens (Kelloniemi *et al*., 2015). **(B)** Formation of a VvLYK1-1 / VvLYK5-1 complex after perception of chitin oligomers at the plasma membrane to activate a strong MAMP-triggered immunity (MTI) in grapevine cells (Roudaire *et al*., 2023). **(C)** Constitutive expression of VvLYK6, which interacts with VvLYK1-1, might deplete VvLYK1-1 for its interaction with VvLYK5-1 and decrease chitin-triggered plant defense responses. Receptors have been classified by colour, VvLYK1-1 = blue, VvLYK5-1 = red, VvLYK6 = green.

During chitin-triggered immunity in *V. vinifera*, *VvLYK6* expression appears to be negatively correlated with the expression of the well-established defense-related gene *VvPAL* and *VvSTS1.2*, encoding the phenylalanine ammonia lyase and the stilbene synthase, respectively. These two enzymes catalyze the final step in the biosynthesis of resveratrol, the main grapevine phytoalexin. When *VvLYK6* is overexpressed in grapevine cells, the expression of *VvPAL* and *VvSTS1.2* is significantly suppressed. These findings are consistent with the observed susceptibility of *V. vinifera* to *B. cinerea* during berry ripening. While green berries typically exhibit basal resistance to *B. cinerea*, mature berries become highly susceptible at harvest. At the veraison stage, Kelloniemi et al. (2015) showed that the defense-related gene *VvPAL* and *VvSTS1.2* are significantly upregulated in green berries infected by *B. cinerea*, leading to the accumulation of resveratrol, while the expression of *VvLYK6* remains very low. In contrast, in mature berries, *VvLYK6* is strongly upregulated during infection, while *VvPAL* and *VvSTS1.2* are not induced, resulting in increased susceptibility to the pathogen (Kelloniemi et al., 2015). Our findings suggest that *VvLYK6* may act as a key molecular component influencing the expression of *VvPAL* and *VvSTS1.2*, thereby modulating grapevine susceptibility to *B. cinerea* infection. Interestingly, similar patterns have been observed in other species, such as tomato, where ripe fruits are significantly more susceptible to *B. cinerea* than unripe ones (Silva et al., 2021). In that study, transcriptomic analyses revealed an upregulation of *SlLYK9*, the tomato ortholog of *VvLYK6* (Figure 1), in ripe fruits infected by the necrotrophic fungi *B. cinerea* and *Rhizopus stolonifer* (Figure S9). As in grapevine, the upregulation of *SlLYK9* correlates with increased susceptibility in mature tomato fruits, suggesting a conserved role in the suppression of plant defense responses (Silva et al., 2021). These observations point to a broader functional role for LysM-RLKs belonging to the LYRI B clade. While LYRI B members have traditionally been associated with the establishment of symbiosis (Li et al., 2022; Ding et al., 2025), emerging evidence suggests they may also act as negative regulators of plant immunity across different organs and plant–fungus interactions. This dual role is supported by their expression in both root and aerial tissues (leaves and fruits) in grapevine and tomato (Figure S10). Such functional versatility has also been reported for other LysM-RLK clades. Members of the LYRI A clade, for example, are involved in symbiosis and can act as either positive or negative regulators of immunity depending on the plant species (Buendia et al., 2018; Zhang et al., 2021), although their expression is typically restricted to roots. Similarly, LYRIII C genes function as negative regulators of both immunity and AM symbiosis and are expressed in both root and aerial tissues (Wang et al., 2023). Taken together, these findings suggest that LYRI B members may serve as repressors of plant immunity in aerial organs during *B. cinerea* infection, while also contributing to AM symbiosis in roots. The potential role of *VvLYK6* in AM establishment warrants further investigation, particularly since, unlike other LYRI B proteins, it does not appear to retain a high affinity for chitooligosaccharides (COS).

From an evolutionary perspective, we analyzed the LYK6 receptor sequence from *Muscadinia rotundifolia (*MrLYK6), a species closely related to *V. vinifera* and naturally resistant to *B. cinerea* (Gabler et al., 2003). Sequence alignment between MrLYK6 and VvLYK6 revealed a high identity of 95.4%, with more than half of the amino acid variations located within the LysM2 domain (Figure S11A). This domain has been shown to play a critical role in chitin binding in OsCEBIP, AtCERK1 and OsCERK1 with specific residues directly involved in ligand interaction (Liu et al.; 2012; Hayafune et al., 2014; Xu et al., 2023). Notably, multiple sequence alignment revealed differences between VvLYK6 and MrLYK6 at several key residues known to be essential for chitin binding in the aforementioned receptors (Figure S11A–B). To further investigate these differences, we used AlphaFold3 to predict the three-dimensional structures of the LysM2 domains of VvLYK6 and MrLYK6, based on the crystallized structure of OsCERK1 bound to a chitin hexamer (Xu et al., 2023). These structural models revealed distinct differences in both electrostatic charge distribution and conformational features within the chitin-binding region (Figure S11C). Taken together, these natural variations in the LysM2 domain of VvLYK6 may explain its lack of detectable affinity for chitin oligomers when expressed alone (Figure 6C). Conversely, the putative chitin-binding capacity of MrLYK6 could contribute to the enhanced resistance of *M. rotundifolia* to *B. cinerea*, highlighting the evolutionary divergence of LysM-RLK function even among closely related species.

To conclude, our study demonstrates that VvLYK6 forms a chitin-induced receptor complex with VvLYK1-1, and that its overexpression negatively regulates plant immune responses, thereby enhancing susceptibility to *B. cinerea* and other fungal pathogens. These findings support the classification of *VvLYK6*, particularly in susceptible *V. vinifera* cultivars, as a susceptibility (*S*) gene. Consequently, targeted knock-out of *VvLYK6* represents a promising strategy to improve grapevine resistance to *B. cinerea* and potentially other fungal pathogens. In line with the growing interest in *S* genes as complementary or alternative targets to classical resistance (*R*) genes in crop improvement (Zaidi et al., 2018), *VvLYK6* emerges as a compelling candidate for genome editing approaches. Furthermore, we identified naturally occurring mutations between *LYK6* from *M. rotundifolia* and *V. vinifera*, which may underlie differences in disease resistance. These beneficial allelic variants could be harnessed through base editing technologies to precisely engineer enhanced resistance traits, offering a novel and sustainable avenue for grapevine breeding.

## Author contributions

JV performed the most of experiments and data analysis. JV, BL, MCH and BP designed the project, experiments and wrote the article. JV produced all transgenic lines of *A. thaliana* excepts lines expressing *VvLYK6* in Col-0 genetic background generated by TR. JV performed all experiments carried out on *A. thaliana* described in this article. NLC and AK transformed and selected grapevine cells for *VvLYK6* overexpression. TM performed all experiments on grapevine cells including treatments and RT-qPCR, Western Blot were done by JV. DL performed transient expression of *VvLYK* in tobacco, microsomal fractions, binding assays and FLIM experiments with CP. TM, CV and VG performed co-IP experiment.

## Acknowledgements

We acknowledge Abdelwahad Echairi and Anna Beslic (SATT Sayens, Dijon, France) for the E. necator strain. We also acknowledge Dr Christian Steinberg for the A. brassicicola inoculum strain MIAE01824 originated from Agroecology unit collection (UMR1347, Dijon, France). We acknowledge Cécile Blanchard for her technical support concerning the infection with the different pathogens. *We acknowledge* Elodie Noirot for confocal microscopy supports from the regional Centre of Microscopy/DImaCell platform (Dijon, France). *We also acknowledge France-BioImaging infrastructure supported by the French National Research Agency (ANR-10-INBS-04)*.

## Supplementary Material

Supplementary Figures S1, S2, S3, S4, S5, S6, S7, S8, S9, S10 and S11 have been uploaded separately in a pdf file.

Supplementary Tables S1, S2, S3 have been uploaded separately in a excel file.

