## Supplementary Figures S1, S2, S3, S4, S5, S6, S7, S8, S9, S10 and S11 for "The *Vitis vinifera* receptor VvLYK6 negatively regulates chitin-triggered immune responses and promotes fungal infections"

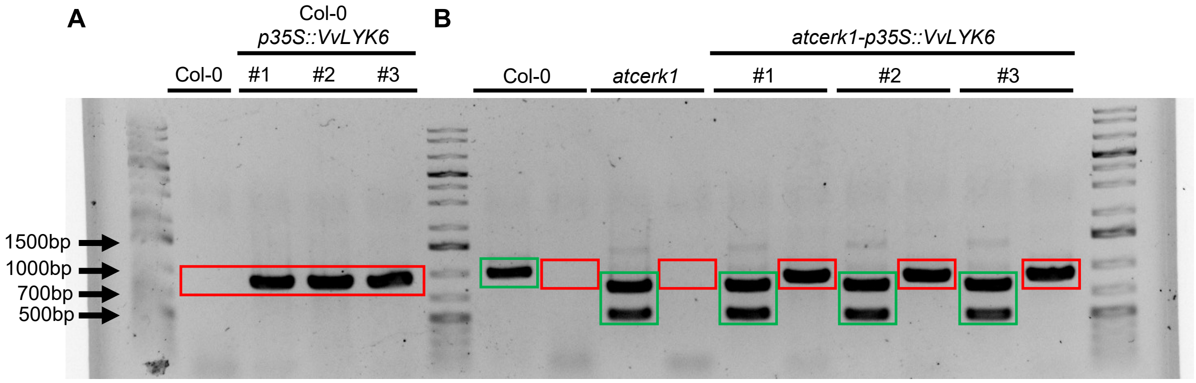


**Figure S1. Genotyping of different lines used in the study. (A-B)** Polymerase chain reaction (PCR) was performed on genomic DNA to check the absence or the presence of the T-DNA in the Col-0 Wild-Type (A) and *Atcerk1* simple mutant (B). Primers used are listed in the supplemental Table S2 and have been designed with the *T-DNA Primer Design* tool of the *SIGnAL* website (http://signal.salk.edu/tdnaprimers.2.html). Expected sizes with the LP + RP primers were between 1000 bp to 1100 bp for the Wild-Type alleles of *AtCerk1* (green). For the T-DNA insertion identification in the simple mutant *atcerk1*, two PCR products with primers BP + RP were obtained at 800 bp and 500 bp for the mutated gene *atcerk1* (green). Presence of *VvLYK6* in all genetic background studied was checked to amplify a region of 867 bp in each transgenic lines and controls (red). Controls were performed on untransformed Wild-Type and the simple mutant *atcerk1*.

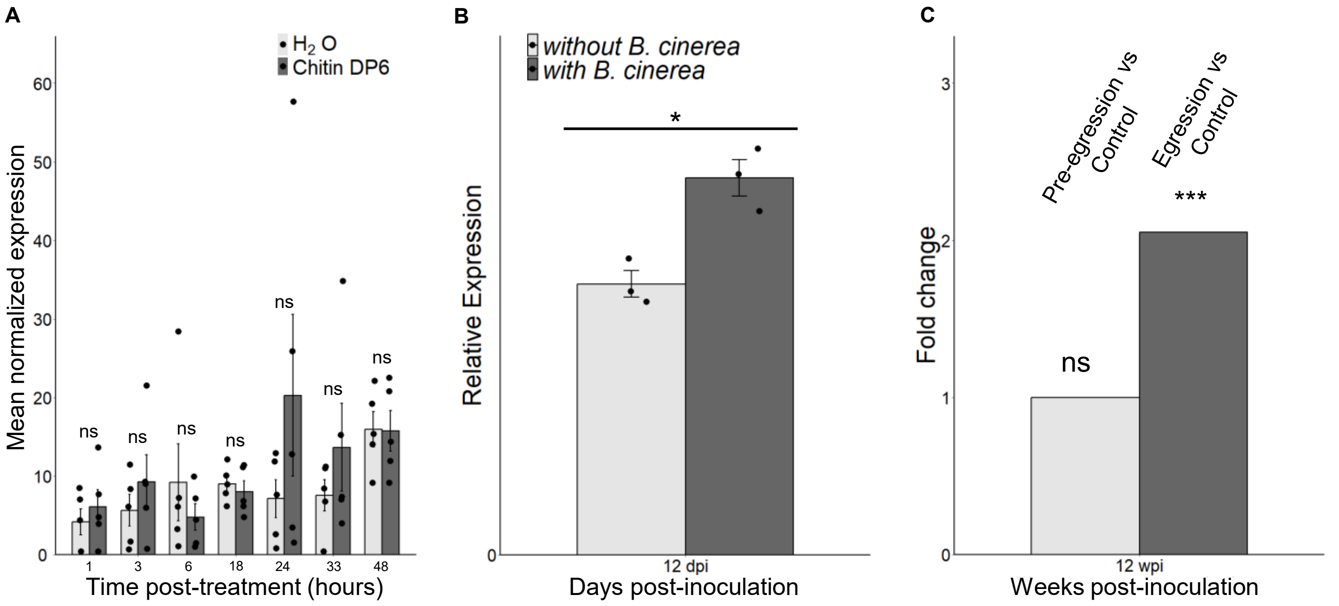


**Figure S2. Supplementary information for *VvLYK6* expression during chitin hexamer treatment and *Botrytis cinerea* infection. (A)** *VvLYK6* expression on grapevine cells 48 h after treatment with chitin DP6. Grapevine cells treated with chitin DP6 were statistically compared to the water-treated cells at each time point with the non-parametric Wilcoxon test (ns, not significant). **(B-C)** *VvLYK6* transcript expression after *B. cinerea* infection in berries from two independent transcriptomic studies. **(B)** *VvLYK6* expression profile analyzed 12 days post-inoculation on mature berries with or without *B. cinerea*. A significant difference between infected and non-infected berries as observed with a non-parametric Kruskal wallis test, *, p-value < 0.05). Using the transcriptomic data from Lovato *et al.* (2019). **(C)** Fold change in *VvLYK6* expression on pre-egression and egression of *B. cinerea* infection versus non-infected ripe berries at 12 weeks post-inoculation. Pre-egression corresponds to quiescent phase of *B. cinerea* infection characterized by the absence of symptoms and egression corresponds to the appearance of symptoms with fungal development. A one-way ANOVA statistical test revealed a significant increase of *VvLYK6* expression only in egressed berries during *B. cinerea* infection (p-value < 0.01 and an absolute fold change ≥ 2.0). Using the transcriptomic data from Haile *et al.* (2020).

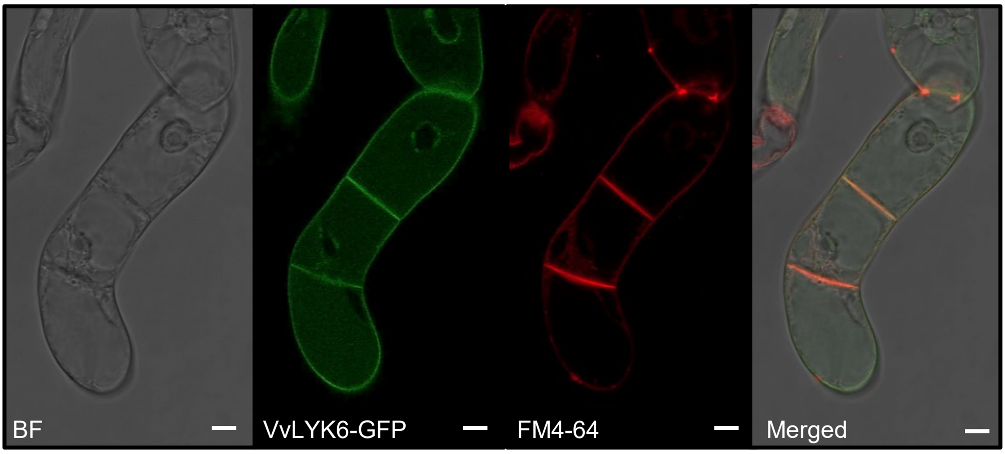


**Figure S3. Subcellular localization of VvLYK6 in grapevine cells cv. Marselan.** GFP-tagged VvLYK6 co-localizes with the plasma membrane marker probe (FM4-64). Scale bars represent 10 µm. Similar localization was observed in two independent lines used for experiments in figure 5. BF = Brightfield.

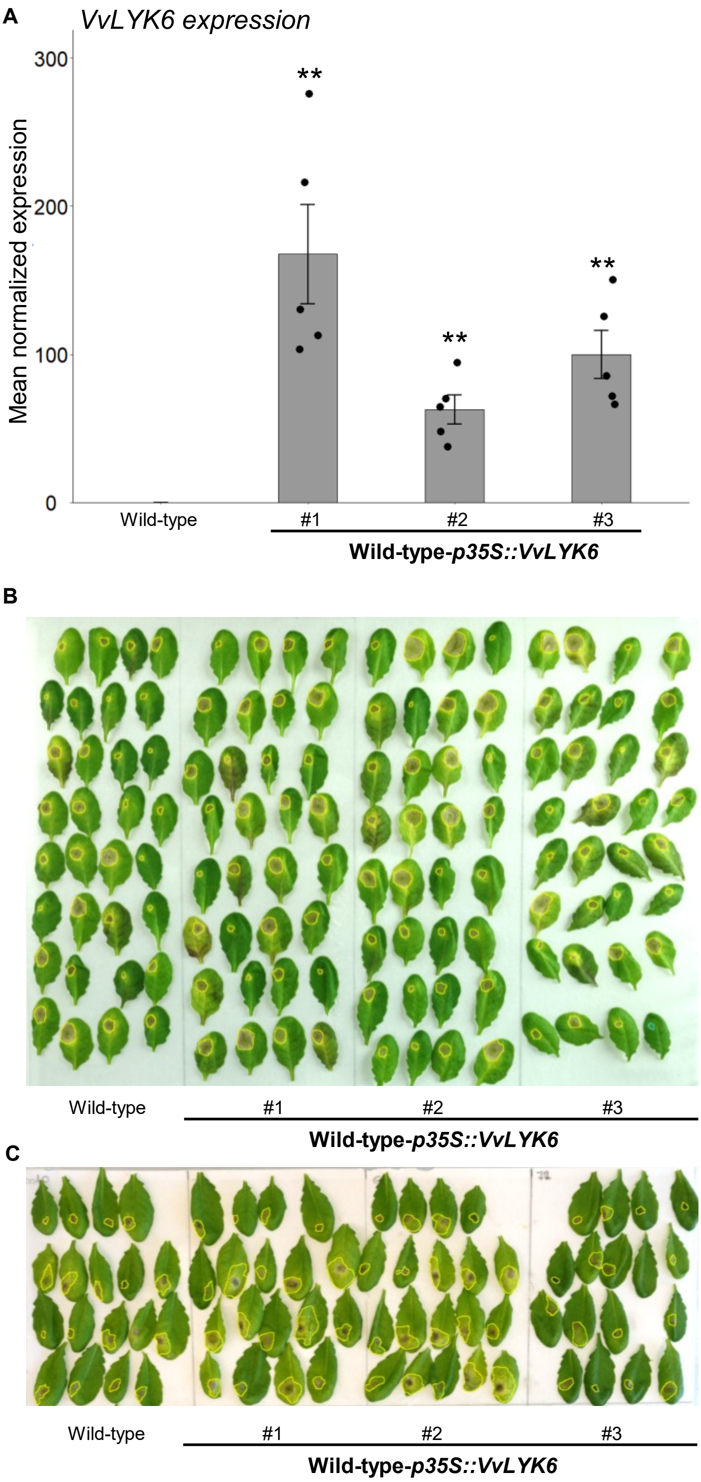


**Figure S4. Relative expression of *VvLYK6* in the three independent lines of *A. thaliana* and representative pathogen assays. (A)** Bars represent the mean of normalized expression level of *VvLYK6* in the three independent lines of *A. thaliana* ± SEM of 5 biologically independent experiments. For each sample, leaves of three different plantlets were sampled. Asterisks indicate a significant difference between independent lines expressing *VvLYK6* and WT (Wilcoxon test, **; P<0.01). The expression levels of *VvLYK6* were normalized to those of two housekeeping genes. **(B)** Representative experiment of Arabidopsis leaves from WT and the three independent lines constitutively expressing *VvLYK6* infected with *B. cinerea* 3 dpi. **(C)** Representative experiment of Arabidopsis leaves from WT and the three independent lines constitutively expressing *VvLYK6* with *A. brassicicola* 5 dpi.

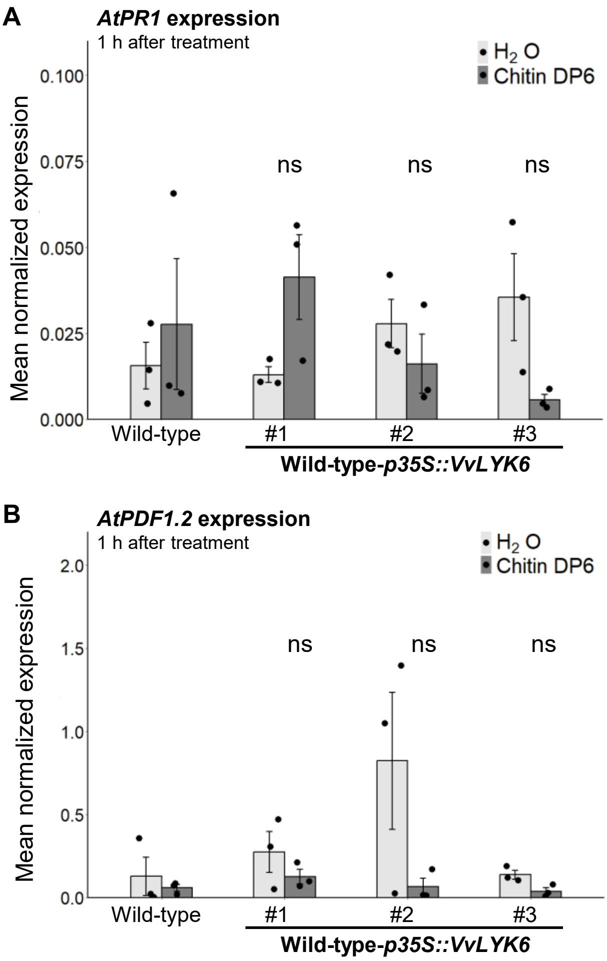


**Figure S5. Gene expression of *AtPR1* and *AtPDF1.2* as marker of salicylic acid (SA) and jasmonic acid (JA) signaling pathways, respectively, in the three independent lines overexpressing *VvLYK6* in *A. thaliana.*** **(A-B)** Normalized expression level of *AtPR1* and *AtPDF1.2* measured by qPCR 1h after chitin DP6 (0.05 g/L) or $H_2O$ treatment. The expression levels of *AtPR1* **(A)** and *AtPDF1.2* **(B)** were normalized to those of two housekeeping genes. Bars represent the mean of relative expression ± SEM of 3 biologically independent experiments. For each sample, leaves of three different plantlets were sampled. A statistical comparison was conducted between the gene expression of the three transgenic lines treated with chitin and the WT treated with chitin (non-parametric Wilcoxon test; ns; no significant).

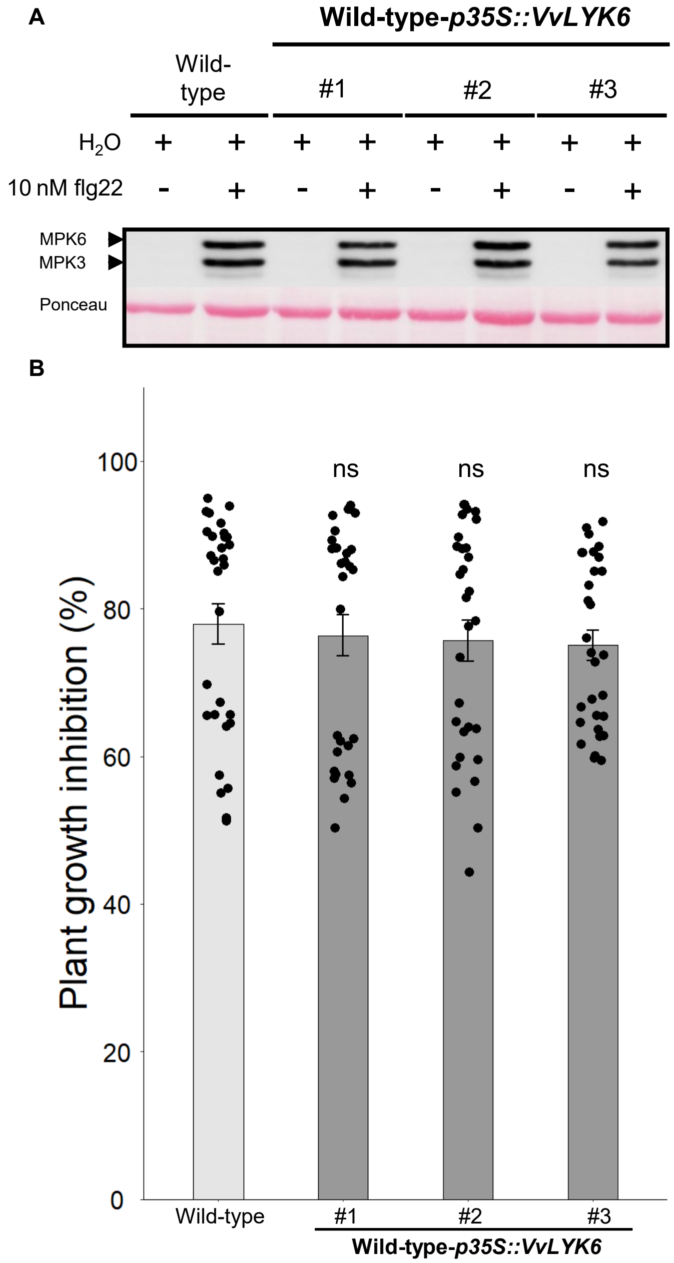


**Figure S6. Constitutive expression of *VvLYK6* has no effect on flg22 elicitation or plant growth inhibition. (A)** Representative immunoblotting with α-pERK1/2 and homogeneous loading verified by Ponceau staining. MAPKs phosphorylation was detected 10 min after $H_2O$ or flg22 (10 nM) treatment by immunoblotting with an antibody α-pERK1/2 on Wild-Type and three independent *VvLYK6*-expressing lines. Similar results were obtained in three biologically independent experiments. For each sample, leaves of three different plantlets were sampled. **(B)** Growth inhibition induced by flg22 is not impacted by the constitutive expression of *VvLYK6* in *A. thaliana*. Five-days old plantlets of *A. thaliana* were grown for 12 days in MS/2 media supplemented with flg22 (1µM) or not. Bars represent the mean growth inhibition (fresh weight of treated/non-treated plantlets) on 30 plantlets from three biologically independent experiments. Non-parametric Wilcoxon test was performed to compare the growth inhibition of each line with the Wild-Type (ns, not significant).

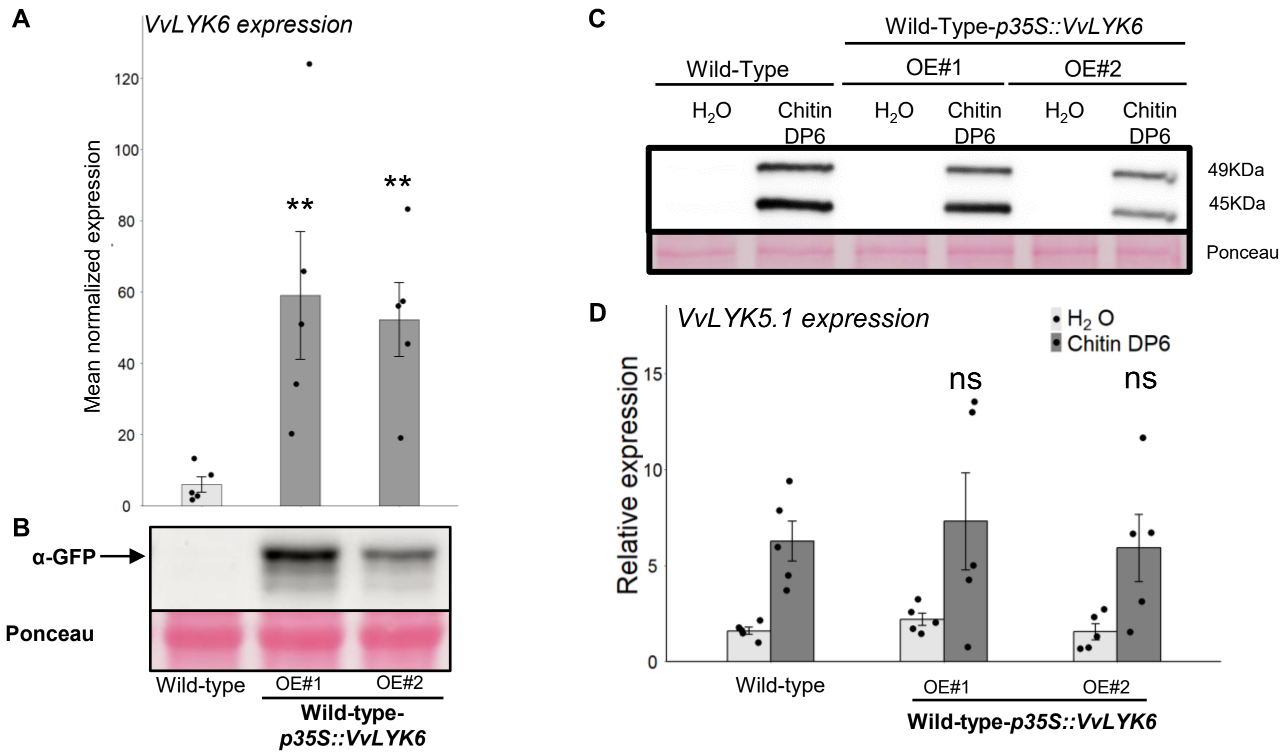


**Figure S7. Expression data and MAPKs phosphorylation and in two independent lines of grapevine cells. (A)** Normalized expression level of *VvLYK6* in the two independent lines of grapevine cells performed by qPCR. The expression levels of *VvLYK6* were normalized to those of two housekeeping genes. Asterisks indicate a significant difference of expression between constitutively expressing *VvLYK6-GFP* transgenic lines and the Wild-Type (Wilcoxon test, **; P<0,01). **(B)** Immunodetection of VvLYK6-GFP accumulation by western blotting in the two transgenic cells overexpressing VvLYK6-GFP and the Wild-Type. **(C)** Representative immunoblotting with α-pERK1/2 of MAPKs activation detected 10 min after $H_2O$ or chitin DP6 treatment (0.05 g/L) in the Wild-Type and two independent grapevine cells lines constitutively expressing *VvLYK6-GFP*. Homogeneous loading was verified by Ponceau red staining. **(D)** Normalized expression level of V*vLYK5.1* measured by qPCR 1h after chitin DP6 (0.05 g/L) or $H_2O$ treatment. The expression levels of *VvLYK5.1* were normalized to those of two housekeeping genes. Bars represent the mean of normalized expression ± SEM of 5 biologically independent experiments. Asterisks indicate a statistically significant difference between transgenic lines and the Wild-Type treated with chitin DP6 using a non-parametric Wilcoxon test (ns, not significant).

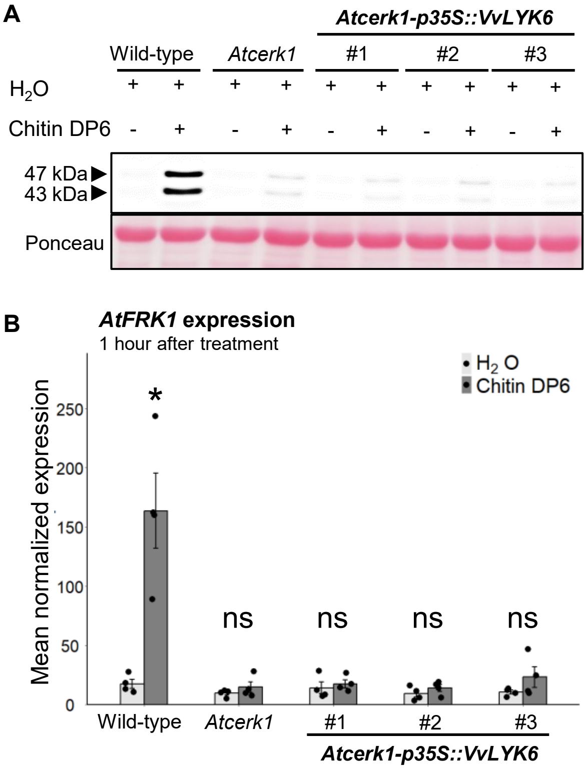


**Figure S8. VvLYK6 does not complement the kinase activity of the *Atcerk1* mutant. (A)** Representative immunoblotting with the pERK1/2 antibody illustrating activation of MAPKs detected 10 min after chitin DP6 (0.05g/L) or $H_2O$ treatment in the WT Col-0, *Atcerk1* and three independent *Atcerk1* lines constitutively expressing *VvLYK6*. Equal protein loading was verified by Ponceau red staining. Similar results were obtained in four biologically independent experiments. **(B)** Normalized expression level of *AtFRK1* measured by qPCR 1h after chitin DP6 (0.05 g/L) or $H_2O$ treatment. Bars represent the mean of normalized expression ± SEM of 4 biologically independent experiments. For each sample, leaves of three different plantlets were sampled. Asterisks indicate a statistically significant difference between chitin DP6 and $H_2O$ condition for each line (Non-parametric Wilcoxon test; *, $P<0.05$; ns, not significant).

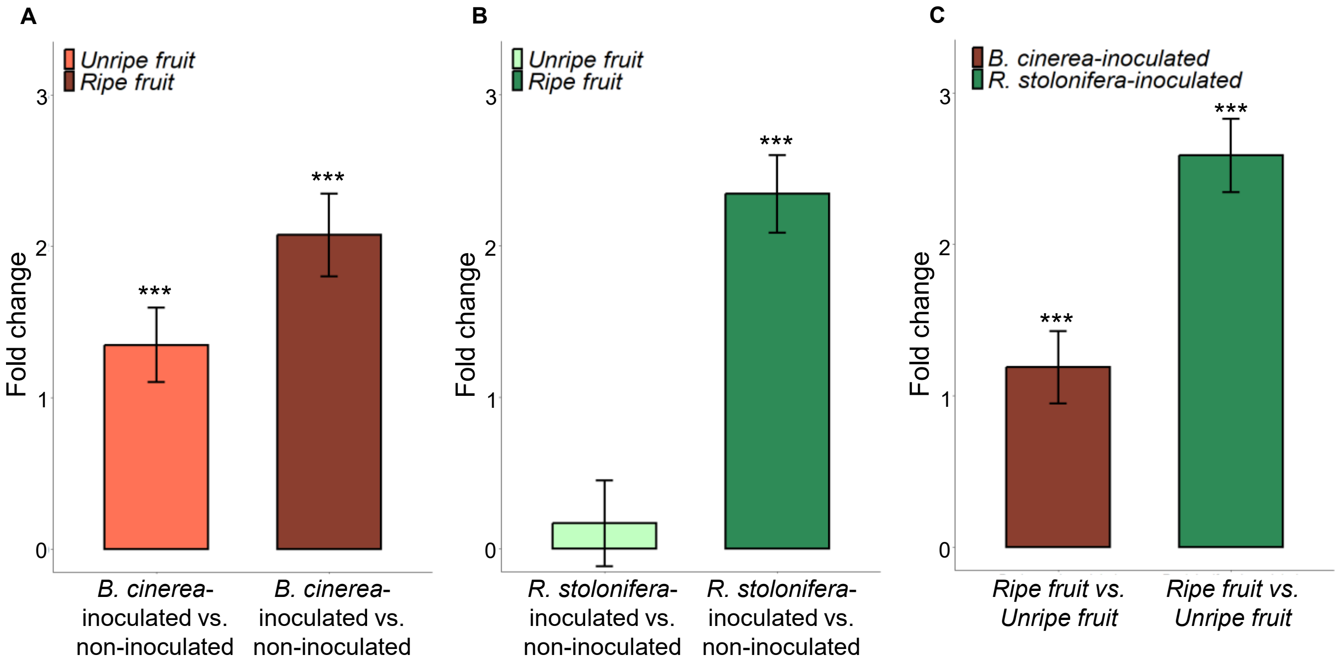


**Figure S9. The *VvLYK6* ortholog *SlLYK9* in *Solanum lycopersicum* (Solyc09g083210) is induced during necrotrophic fungi *Botrytis cinerea* and *Rhizopus stolonifera* infection in ripe tomato fruit. (A)** Fold change of *SlLYK9* expression on *B. cinerea* inoculated versus non-inoculated unripe and ripe tomato. **(B)** Fold change of *SlLYK9* expression on *R. stolonifera* inoculated versus non-inoculated unripe and ripe tomato. **(C)** Fold change of *SlLYK9* expression during *B. cinerea* and *R. stolonifera* infection on inoculated ripe versus inoculated unripe tomato. All transcriptomic data correspond to one day post inoculation and are from Silva *et al.* (2021). A statically Wald test were performed for each comparison (***, p-value < 0.001).

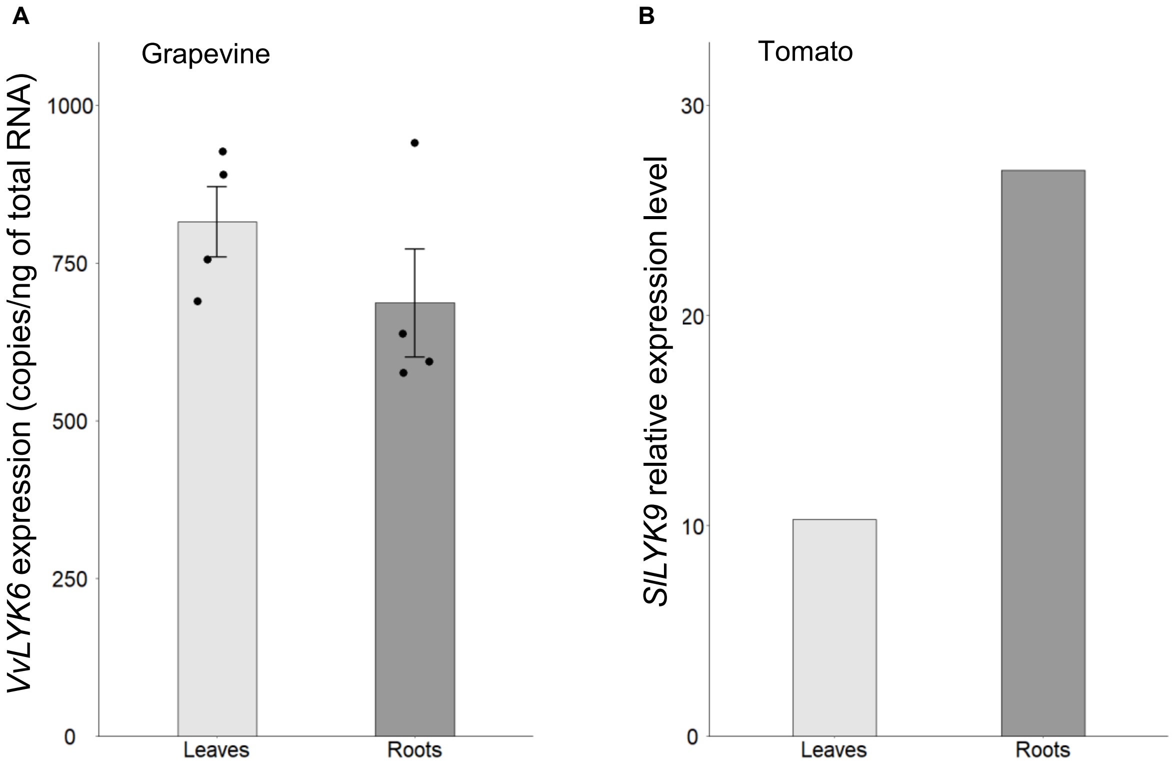


**Figure S10. Expression of *VvLYK6* and *SlLYK9* orthologs in leaves and roots of grapevine and tomato respectively. (A)** *VvLYK6* transcript levels in grapevine (*V. vinifera* cv. Chardonnay) leaves and roots from *in vitro* plantlets. RT-qPCR was performed on first-strand cDNAs synthesize from total RNAs. **(B)** Expression level of *SlLYK9* in tomato (*S. lycopersicum* cv. Heinz) leaves and roots. The data were taken from TomExpress (Zouine et al., 2017).

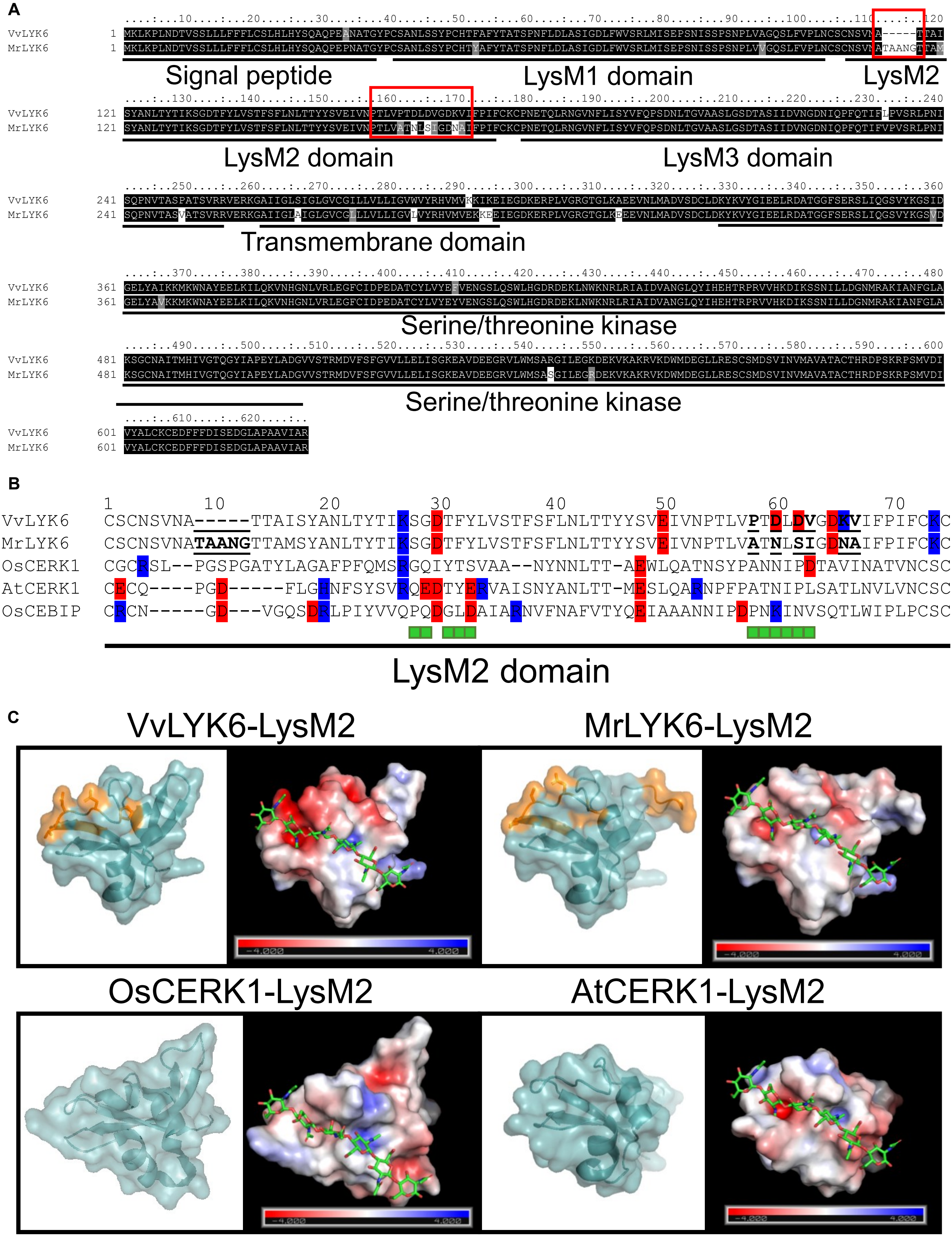
A
VvLYK6
MrLYK6
Signal peptide
LysM1 domain
LysM2
LysM2 domain
LysM3 domain
Transmembrane domain
Serine/threonine kinase
Serine/threonine kinase
B
VvLYK6
MrLYK6
OsCERK1
AtCERK1
OsCEBIP
LysM2 domain
C
VvLYK6-LysM2
MrLYK6-LysM2
OsCERK1-LysM2
AtCERK1-LysM2

**Figure S11. Identification of mutations in the LysM2 domain between LYK6 from the susceptible species *Vitis vinifera* cv. Marselan (VvLYK6) and the resistant species *Muscadinia rotundifolia* (MrLYK6). (A)** Clustal alignment of VvLYK6 and MrLYK6 performed with MEGAX (Kumar *et al.* 2018). Amino acids conservation is highlighted with boxshade colours with black corresponding to conserved residues and white corresponding to divergent residues. Known domains of LysM receptor-like kinase are underlined with a black line including signal peptide, LysM1, LysM2, LysM3, transmembrane and serine/threonine kinase. Region with several mutations are highlighted with red boxes. **(B)** Clustal alignment of LysM2 domain from VvLYK6, MrLYK6 and three crystallized LysM2 domains from OsCERK1, OsCEBIP and AtCERK1 (Liu *et al.*, 2012; Liu *et al.,* 2016; Xu *et al.*, 2023) performed with MEGAX. Mutations between VvLYK6 and MrLYK6 are underlined in bold. Positively and negatively charged residues are highlighted in blue and red respectively. Green squares indicate the residues of AtCERK1 and OsCERK1 involved in chitin binding. **(C)** The predicted structure of the LysM2 domain of VvLYK6 and MrLYK6 was generated using AlphaFold 3 and visualized with PyMOL software, based on the crystallized LysM2 of OsCERK1. On the left, the putative three-dimensional structures of each protein are shown with a cyan surface at 40% transparency. Mutated residues between VvLYK6 and MrLYK6 are highlighted in orange using stick representation. On the right, the chitin hexamer binding site on the surface of LysM2 is depicted, based on the crystallized chitin-associated LysM2 of OsCERK1 (Xu *et al.*, 2023). The LysM2 domain is displayed using an electrostatic surface potential map, where blue, white, and red represent positive, neutral, and negative surface charges, respectively.
